# Functional Interrogation of *HOXA9* Regulome in MLLr Leukemia via Reporter-based CRISPR/Cas9 screen

**DOI:** 10.1101/2020.04.20.050583

**Authors:** Hao Zhang, Yang Zhang, Shaela Wright, Judith Hyle, Lianzhong Zhao, Jie An, Xinyue Zhou, Xujie Zhao, Ying Shao, Hyeong-Min Lee, Taosheng Chen, Yang Zhou, Rui Lu, Chunliang Li

## Abstract

Aberrant *HOXA9* expression is a hallmark of most aggressive acute leukemias, including human acute myeloid leukemia (AML) and subtypes of acute lymphoblastic leukemia (ALL). *HOXA9* overexpression not only predicts poor diagnosis and outcome but also plays a critical role in leukemia transformation and maintenance. However, our current understanding of *HOXA9* regulation in leukemia is limited, hindering development of therapeutic strategies to treat *HOXA9*-driven leukemia. To mitigate these challenges, we generated the first *HOXA9-mCherry* knock-in reporter in an MLL-rearranged (MLLr) B-ALL cell line to dissect *HOXA9* regulation. By utilizing the reporter and CRISPR/Cas9 mediated screens, we identified transcription factors controlling *HOXA9* expression, including a novel regulator, USF2 and its homolog USF1. USF1/USF2 depletion significantly down-regulated *HOXA9* expression and impaired MLLr leukemia cell proliferation. Ectopic expression of HOXA9-MEIS1 fusion protein rescued the impaired leukemia cell proliferation upon USF2 loss. Cut&Run analysis revealed the direct occupancy of USF2 onto *HOXA9* promoter in MLLr leukemia cells. Collectively, the *HOXA9* reporter facilitated the functional interrogation of the *HOXA9* regulome and has advanced our understanding of the molecular regulation network in *HOXA9*-driven leukemia.

## INTRODUCTION

Dysregulation of the homeobox (HOX)-containing transcription factor *HOXA9* is a prominent feature in most aggressive acute leukemias (1, 2). During normal hematopoiesis, HOXA9 plays a critical role in hematopoietic stem cell expansion and is epigenetically silenced during lineage differentiation (2). In certain leukemia subtypes, this regulatory switch fails and *HOXA9* is maintained at high levels to promote leukemogenesis. However, the mechanisms governing *HOXA9* expression remain to be fully understood. *HOXA9* overexpression is commonly observed in over 70% of human acute myeloid leukemia (AML) cases and ~10% of acute lymphoblastic leukemia (ALL) cases (3). Notably, the high expression of *HOXA9* is sharply correlated with poor prognosis and outcome in human leukemia (4, 5). An accumulating body of evidence indicates that *HOXA9* dysregulation is both sufficient and necessary for leukemic transformation (1, 2). Forced expression of *HOXA9* enforces self-renewal, impairs myeloid differentiation of murine marrow progenitors, and ultimately leads to late onset of leukemia transformation (6), which is accelerated by co-expression with interacting partner protein MEIS1 (7). Conversely, knocking down *HOXA9* expression results in leukemic cell differentiation and apoptosis (8, 9). Thus, excessive *HOXA9* expression has emerged as a critical mechanism of leukemia transformation in many hematopoietic malignancies.

Consistent with the broad overexpression pattern of *HOXA9* in many leukemia cases, a wide variety of genetic alterations in leukemia contribute to *HOXA9* dysregulation, including *MLL* gene rearrangements (MLLr), *NPM1* mutations, *NUP98*-fusions, *EZH2* loss-of-function mutations, *ASXL1* mutations, *MOZ* fusions and other chromosome alterations (1, 3, 10, 11). Additionally, our recent work shows that *DNMT3A* hotspot mutations may also contribute to *HOXA9* overexpression by preventing DNA methylation at its regulatory regions (12). Given that genomic variation of *HOXA9* including NUP98-HOXA9 fusion and gene amplification accounted for less than 2% of *HOXA9* overexpression in AML cases (13–15), uncovering the upstream epigenetic and transcriptional regulators of *HOXA9* in leukemia could advance the design of novel therapeutic interventions. For example, because MLLr proteins recruit the histone methyltransferase DOT1L to the *HOXA* locus, promoting hyper-methylation at histone H3 lysine 79 and subsequent high *HOXA9* transcription (16), selective DOT1L inhibitors have been exploited to inhibit leukemia development and *HOXA9* expression in MLLr leukemias and are now in clinical trials (17, 18). However, DOT1L inhibitors usually act slowly and their effects remain sub-optimal. Remarkably, drug resistance was inevitably observed (19, 20), suggesting that other regulators may be involved in sustaining aberrant *HOXA9* expression. To date, most known *HOXA9* regulator proteins are epigenetic modifiers, and little is known about which DNA-binding transcription factors are involved in directly regulating *HOXA9* expression in acute leukemia (19, 21–23).

Previous studies have also advocated that the organization of chromatin domains at the *HOXA* gene cluster contributes to high *HOXA9* expression in cancer cells (24, 25). Specifically, CCCTC-binding factor CTCF may potentiate *HOXA9* expression through direct binding at the conserved motif between *HOXA7* and *HOXA9* (CBS7/9) to establish necessary chromatin looping interaction networks in AML (24). In contrast, Ghasemi et al. (BioRxiv, https://doi.org/10.1101/2020.02.17.952390) reported that *HOXA* gene expression was maintained in the CTCF binding site deletion mutants, suggesting that transcriptional activity at the *HOXA* locus in NPM1-mutant AML cells does not require long-range CTCF-mediated chromatin interactions. However, whether loss of CTCF has a direct effect on *HOXA9* expression remains to be studied. Lastly, although the clinical significance of *HOXA9* has been recognized for more than two decades, it is technically difficult to systematically discover regulators of *HOXA9* in acute leukemia owing to the lack of an endogenous reporter to dictate *HOXA9* expression.

In this work, we sought to establish an endogenous reporter system, enabling real-time monitoring of *HOXA9* expression in conjunction with high-throughput CRISPR/Cas9 screening in a human B-ALL MLL-rearranged t(4,11) cell line, SEM, equipped with an endogenous *HOXA9^P2A-mCherry^* reporter allele. The *HOXA9^P2A-mCherry^* reporter allele authentically recapitulated endogenous transcription of the *HOXA9* gene and did not affect endogenous transcription of other adjacent *HOXA* genes. To gain a global understanding of the transcription factors regulating *HOXA9* expression, we performed a CRISPR/Cas9 loss-of-function screen specifically targeting 1,639 human transcription factors. Our screening robustly re-identified expected targets such as *DOT1L* and *HOXA9* itself. More importantly, we identified novel functional regulators of *HOXA9* including Upstream Transcription Factor 2 (USF2). Surprisingly, the CRISPR screen and global depletion of CTCF via siRNA and degron-associated protein degradation all demonstrated that *HOXA9* does not down-regulate upon CTCF loss. We conclude that the *HOXA9^P2A-mCherry^* reporter serves as a robust tool for discovery of novel *HOXA9* regulators.

## RESULTS

### Establishment and characterization of the*HOXA9^P2A-mCherry^* reporter human MLLr leukemia cell line

As shown by many previous studies, *HOXA9* overexpression was observed in refractory MLL-rearranged ALL and AML patients (26–28)(Figures S1A-S1C). Therefore, we utilized our previously reported high-efficiency knock-in strategy, “CHASE knock-in” (29), to deliver the *P2A-mCherry* cassette upstream of the *HOXA9* stop codon in a patient-derived human B-ALL cell line, SEM, which has a typical B-ALL signature along with a t(4;11) translocation and maintains one single allele of the *HOXA* gene cluster. Because the P2A-mediated ribosome skipping disrupts the synthesis of the glycyl-prolyl peptide bond at the C-terminus of the P2A peptide, translation leads to dissociation of the P2A peptide and its immediate downstream mCherry protein (30). Therefore, the knock-in allele would produce a functional HOXA9 protein under control of the endogenous promoter and intrinsic *cis*-regulatory elements while delivering a separate mCherry protein. In brief, we constructed the knock-in vector containing a *P2A-mCherry* cassette flanked with 5’ and 3’ *HOXA9* homology arms (HAs) of approximately 800-bps, which were cloned from SEM cells. A single guide RNA (sgRNA) and a protospacer adjacent motif (PAM) sequence targeting the genomic sequence 5’ of the *HOXA9* stop codon was inserted into the 5’ ends of both HAs (Figure 1A). When the HA/knock-in cassette was co-electroporated with an all-in-one vector expressing wild-type Cas9 and the same *HOXA9* sgRNA, the HA/knock-in cassette was released from the donor vector with two nuclease cleavages and delivered to the target genomic region where double-strand breaks occurred. Successful knock-in cells were enriched by flow cytometry sorting for mCherry (Figure 1B) and characterized via genotyping PCR and Sanger sequencing (Figure 1C). To examine the possibility of random integration of the *P2A-mCherry* cassette, fluorescence *in situ* hybridization (FISH) was performed with a *P2A-mCherry* DNA probe (red) and a FITC-labeled fosmid DNA probe targeting the *HOXA9* locus (green). On-target knock-in cells displayed co-localization of red and green fluorescence without random integration signals in the rest of genome (Figure 1D and Figures S2A-S2D). The bulk knock-in population, hereafter called *HOXA9^P2A-mCherry^*, was used as a reporter cell line for the entire study. Many knock-in studies reported the exogenous DNA fragment may affect normal endogenous gene expression in a complex chromatin niche (31, 32). Therefore, to test whether the inserted *P2A-mCherry* segment would affect the gene expression pattern of *HOXA9* and its neighboring *HOXA* cluster genes, Q-PCR analysis was conducted on both wild-type (WT) and *HOXA9^P2A-mCherry^* knock-in (KI) cells. RNA-seq data collected from SEM cells in our previous studies suggested that *HOXA7*, *HOXA9* and *HOXA10* were the only highly expressed *HOXA* genes in MLLr leukemia SEM cells (29)(Figure 1E), and that these patterns were indistinguishable between WT and KI populations, indicating the *P2A-mCherry* knock-in did not alter the gene expression landscape at the *HOXA* cluster (Figure 1F).

**Figure 1.**
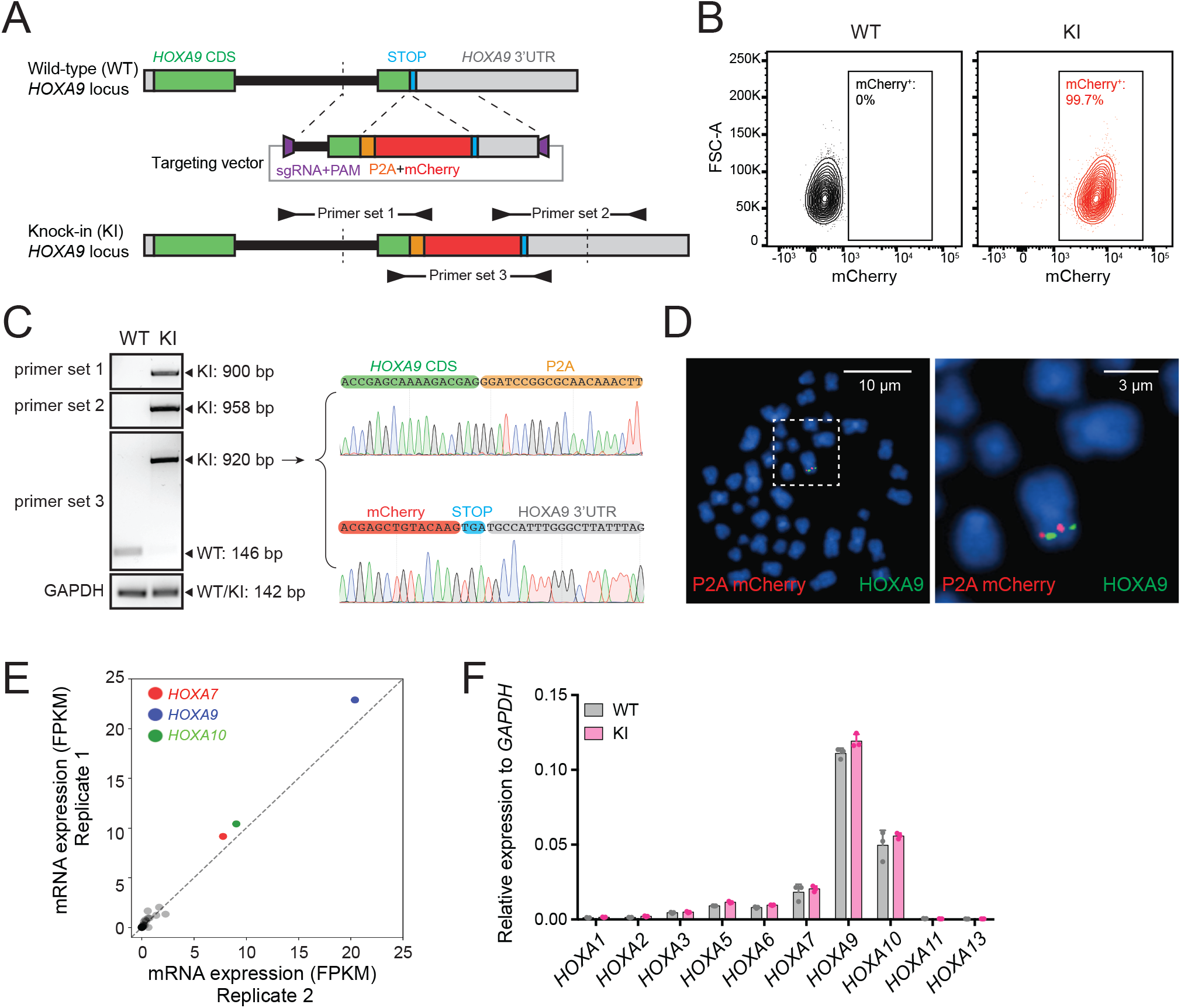
Establishment and characterization of the *HOXA9^P2A-mCherry^* reporter human MLLr leukemia cell line. (A) Schematic diagram of the knock-in design and genotyping PCR primer design for the *HOXA9^P2A-mCherry^* reporter allele. (B) Flow cytometry analysis of *HOXA9^P2A-mCherry^* reporter cells. Wild-type SEM cells were used as negative controls. (C) Genotyping PCR products from the 5′ and 3′ knock-in boundaries were sequenced to verify the seamless knock-in of the *mCherry* reporter gene to the endogenous locus. (D) Fluorescence *in situ* hybridization of the *P2A-mCherry* knock-in cassette in *HOXA9^P2A-mCherry^* reporter cells. The *P2A-mCherry* DNA was labeled with a red-dUTP by nick translation, and an *HOXA9 BAC* clone was labeled with a green-dUTP. The cells were then stained with 4,6-diamidino-2-phenylindole (DAPI) to visualize the nuclei. A representative metaphase cell image is shown for the pattern of hybridization (pairing of red and green signals). (E) RNA-seq data of all *HOXA* cluster genes were illustrated as log_2_ (normalized numbers of FPKM) from two replicate samples of SEM cells. *HOXA7*, *HOXA9* and *HOXA10* were highlighted by color code. (F) Q-PCR analysis confirmed the unaffected *HOXA* cluster gene transcription between *HOXA9^P2A-mCherry^* reporter (KI) and WT SEM cells. Data shown are means ± SEM from replicate independent experiments. *p < 0.05 of two-tailed Student’s t test.

### The *HOXA9^P2A-mCherry^* reporter allele recapitulates endogenous transcription of *HOXA9* in MLLr SEM cells

To evaluate whether the *HOXA9^P2A-mCherry^* reporter allele would faithfully respond to the transcriptional regulation of the cellular *HOXA9* promoter, we genetically perturbed or pharmaceutically inhibited *HOXA9*’s upstream regulators. Previous studies have shown that *DOT1L* and *ENL* positively regulate *HOXA9* expression in MLLr leukemia via direct occupancy on *HOXA9*’s promoter (9, 17). Therefore, two sgRNAs targeting the coding region of *DOT1L* (sgDOT1L) and *ENL* (sgENL) were infected into the *HOXA9^P2A-mCherry^* cells expressing Cas9. Flow cytometry and Q-PCR analysis each revealed that *mCherry* and *HOXA9* expression were both down-regulated by sgRNAs targeting *DOT1L* or *ENL* (Figures 2A-2D), and that the mCherry expression correlated well with the expression of HOXA9 (Figure 2E). Additionally, a DOT1L-selective inhibitor, SGC0946 (22), was supplemented at different dosages for 6 days to the *HOXA9^P2A-mCherry^* cells in culture resulting in a dosage-dependent reduction of mCherry fluorescence intensity measured by fluorescence imaging (Figures 2F-2G) and flow cytometry (Figure 2H). Again, Q-PCR analysis of the DMSO- and SGC0946-treated cells showed that mRNA expression of *mCherry* was significantly correlated with that of *HOXA9* (Pearson’s *r*=0.90, p<0.001) (Figure 2I). Taken together, these data confirm that the newly established *HOXA9^P2A-mCherry^* allele was authentically controlled by the endogenous *HOXA9* promoter and its local chromatin niche.

**Figure 2.**
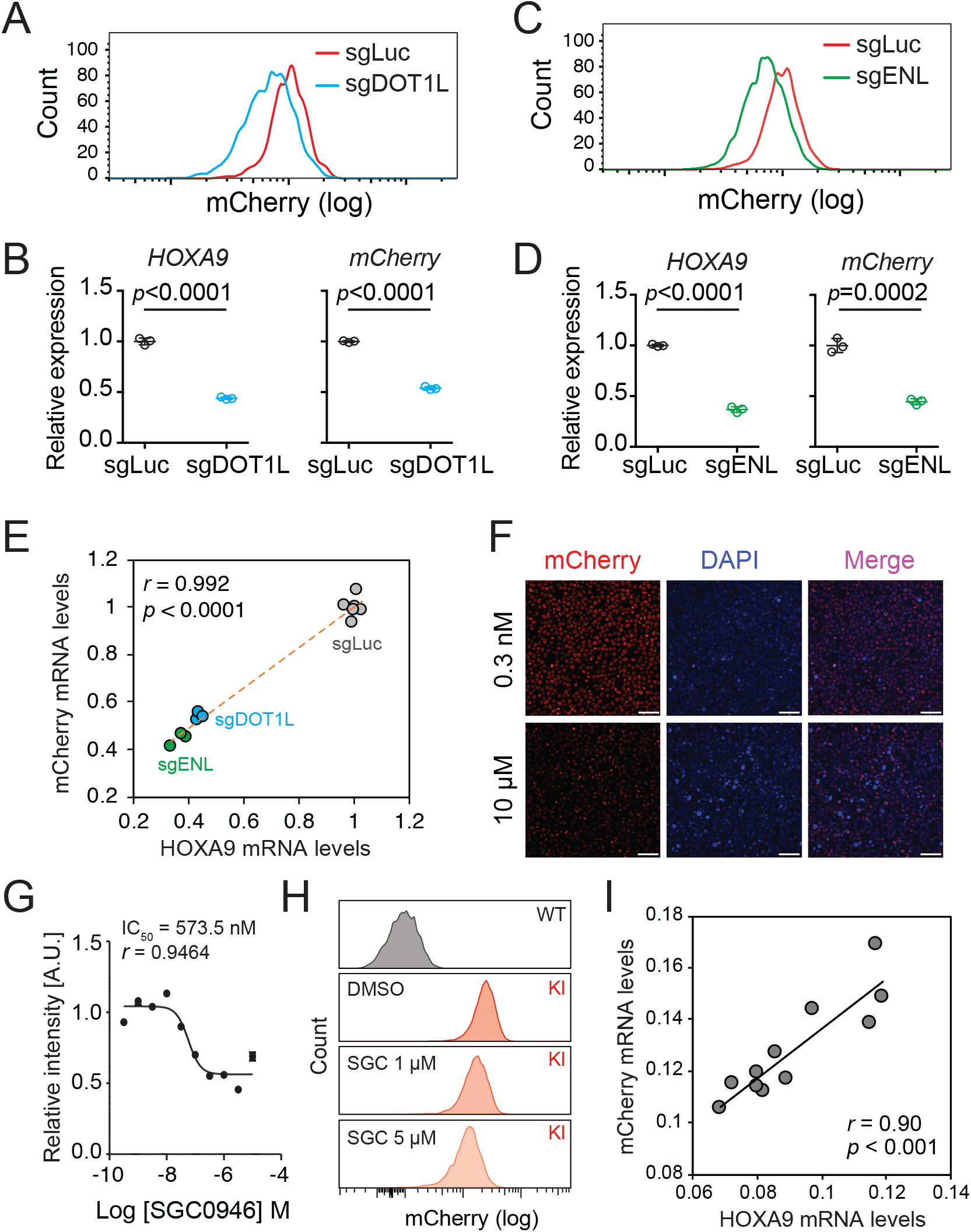
The *HOXA9^P2A-mCherry^* reporter allele recapitulates endogenous transcription of *HOXA9* in MLLr SEM cells. (A) Flow cytometry analysis of the *HOXA9^P2A-mCherry^* cells targeted with luciferase-sgRNA and DOT1L-sgRNA. (B) Q-PCR analysis of the *HOXA9^P2A-mCherry^* cells targeted with luciferase-sgRNA and DOT1L-sgRNA by using specific primers targeting the mRNA sequences of *mCherry* and *HOXA9*. Three biological replicates were performed. Data shown are means ± SEM from replicate independent experiments. The *P*-value was calculated by performing a two-tailed *t*-test. (C) Flow cytometry analysis of the *HOXA9^P2A-mCherry^* cells targeted with luciferase-sgRNA and ENL-sgRNA. (D) Q-PCR analysis of the *HOXA9^P2A-mCherry^* cells targeted with luciferase-sgRNA and ENL-sgRNA by using specific primers targeting the mRNA sequence of *mCherry* and *HOXA9*. Three biological replicates were performed. The *P*-value was calculated by performing a two-tailed *t*-test. (E) The correlation of transcription reduction in *mCherry* and *HOXA9* in response to CRISPR–mediated targeting was calculated by Pearson’s correlation test. (F) Fluorescence imaging was performed on the *HOXA9^P2A-mCherry^* cells treated with various dosages of DOT1L inhibitor SGC0946 for six days. Representative images were shown for comparison between 0.3 nM and 10 μM dosages. For each dosage treatment, four replicates were conducted (scale bar 50 μm). (G) Fluorescence curve was generated according to mCherry intensity in response to dosage dependent treatment of drug for six days. About 20,000 cells were split in each of the 384-well at the starting time point. (H) Flow cytometry analysis of the *HOXA9^P2A-mCherry^* cells treated with DMSO and various dosages of the DOT1L inhibitor SGC0946. (I) Q-PCR analysis of the *HOXA9^P2A-mCherry^* cells with or without the six-day treatment of the DOT1L inhibitor SGC0946 by using specific primers targeting the mRNA sequences of *mCherry* and *HOXA9*. The correlation of transcription reduction in *mCherry* and *HOXA9* in response to inhibitor–mediated transcription repression was calculated by performing Pearson’s correlation test.

### Pooled CRISPR/Cas9 screening identified a novel transcription factor, USF2 that regulates *HOXA9* expression

Although a few regulators of *HOXA9* in MLLr leukemia have been previously identified (9, 11, 33–39), to date a comprehensive CRISPR/Cas9 screen to unbiasedly identify novel upstream regulatory factors of *HOXA9* has not been feasible owing to the lack of a reliable reporter cell line. Therefore, we combined the *HOXA9^P2A-mCherry^* reporter line and an in-house CRISPR-Cas9 sgRNA library targeting 1,639 human transcription factors to identify novel regulatory effectors (40). In this library, seven sgRNAs spanning multiple coding exons were designed per transcription factor, seven sgRNAs targeting DOT1L were included as a positive control, and an additional 100 non-targeting sgRNAs were included as negative controls. Two paralleled screens were performed on the same *HOXA9^P2A-mCherry^* reporter line stably expressing Cas9 and the lentiviral sgRNA library at a low M.O.I. (less than 0.3). Cells were selected with antibiotics, enriched, and fractionated by flow cytometric sorting for the top 10% (mCherry^High^) and bottom 10% (mCherry^Low^) mCherry populations, followed by genomic DNA extraction, PCR, and deep sequencing to identify differentially represented sgRNAs (Figure 3A). Pearson’s correlation implied that global sgRNA distribution significantly differed in the sorted populations of mCherry^High^ and mCherry^Low^ and was similarly correlated between biological replicates (Figure S3A). The differentially represented sgRNAs were calculated by DEseq2 analysis and combined for MAGeCK testing at the gene level (41). In the gene ranking list based on fold-change enrichment of sgRNAs between mCherry^High^ and mCherry^Low^ populations, the top and third-top hits are *HOXA9* and *DOT1L*, suggesting that the screening was successful. In addition, the second-top hit, USF2, was significantly enriched as a novel positive regulator of *HOXA9* (Figure 3B). Consistent with the significant enrichment of these three candidates at the gene level, DEseq2 analysis (42) and sgRNA enrichment plotting both suggested that most of the sgRNAs against these genes were differentially represented (Figures 3C-3D and Figures S3C). Importantly, all of the non-targeting control sgRNAs were similarly distributed across mCherry^High^ and mCherry^Low^ populations, indicating that the sorting-based screen did not bias the enrichment.

**Figure 3.**
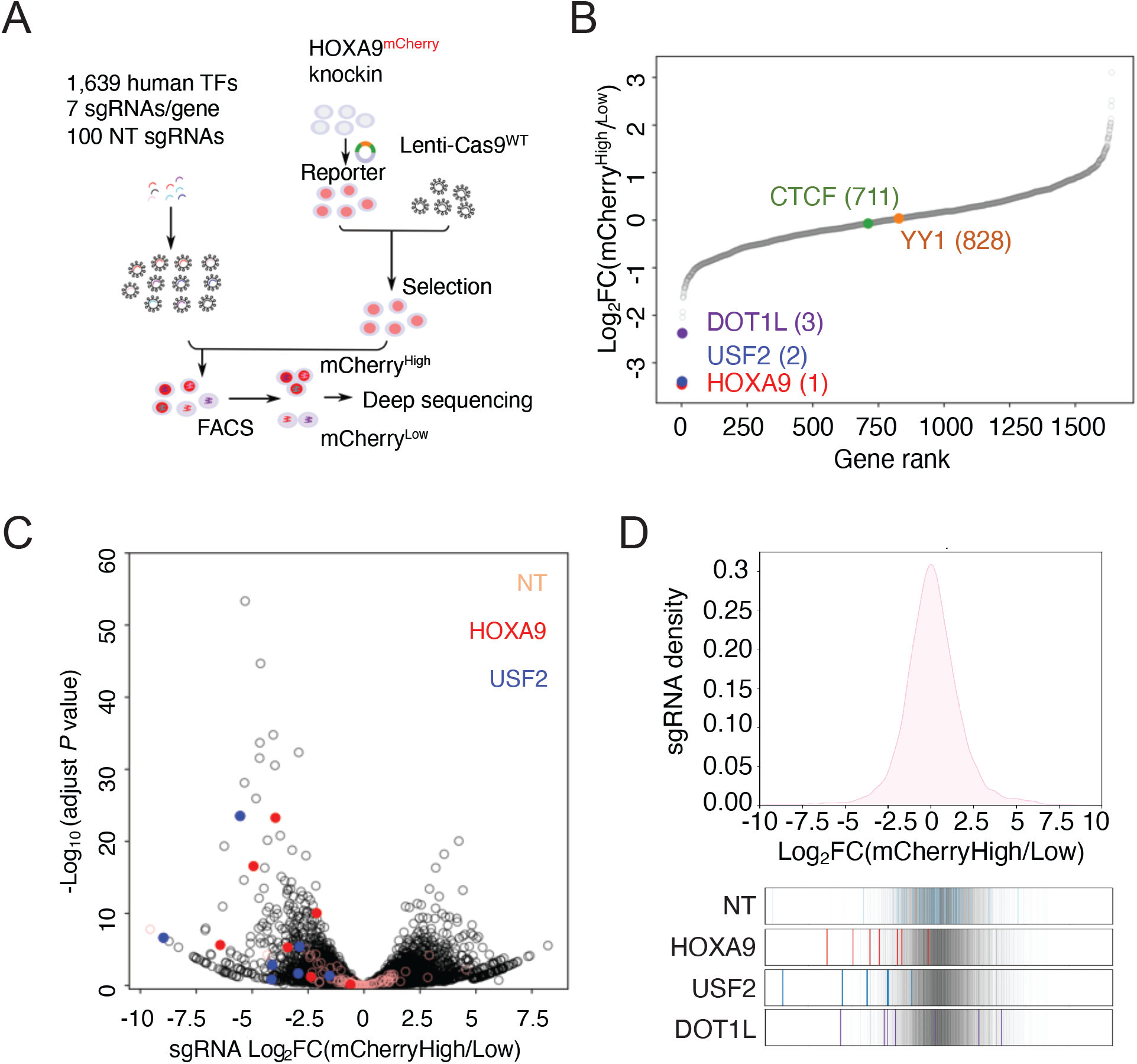
Pooled CRISPR/Cas9 screening identified a novel transcription factor USF2 regulating *HOXA9*. (A) Schematic diagram of a working model of loss-of-function CRISPR screening targeting 1,639 human transcription factors. (B) Gene ranking of all transcription factors from screening was illustrated with *HOXA9*, *USF2*, *DOT1L*, *CTCF* and *YY1* highlighted. The enrichment score of seven sgRNAs against each transcription factor was combined by the MAGeCK algorithm. (C) Top: the overall distribution of all sgRNAs from the screening was shown based on the p-value and the DEseq2 score calculated by Log_2_[Fold Change (mCherry^High^/mCherry^Low^)]. Bottom: NT, *HOXA9* and *USF2* sgRNAs were highlighted by different color code. (D) The ratio for all sgRNAs targeting the top 2 hits, *HOXA9* and *USF2*, are shown between mCherry^High^ and mCherry^Low^ sorted population. NT sgRNAs were overlaid on a gray gradient depicting the overall distribution. NT: 100 sgRNAs. Transcription factors: 7 sgRNAs/each.

### CTCF is dispensable for maintaining *HOXA9* expression in MLLr SEM cells

Interestingly, the most-characterized looping factors, CTCF and YY1, were not enriched in the *HOXA9^P2A-mCherry^* reporter screen (Figure 3B). CTCF was reported to be essential for *HOXA9* expression by occupying the boundary sequence between *HOXA7* and *HOXA9* (CBS7/9) in MLLr AML cell line MOLM13 (24). CRISPR-mediated deletion of the core sequence CTCF binding motif in CBS7/9 significantly decreased *HOXA9* expression and tumor progression (24, 43). Given that CTCF is generally essential for cell survival, it is possible that cells targeted by CTCF sgRNAs in the *HOXA9^P2A-mCherry^* reporter and TF screen quickly dropped out of the population and were unable to be enriched as a regulator of *HOXA9*. To mitigate the challenge, we utilized a previously described auxin-inducible degron (AID) cellular system (29, 44–46) to acutely deplete the CTCF protein in SEM cells and evaluate the immediate transcriptional response of *HOXA9* (Figure S4A). Upon acute depletion of CTCF via auxin (IAA) treatment in three CTCF^AID^ bi-allelic knock-in clones, the protein expression of a previously identified vulnerable gene, *MYC*, was significantly inhibited while HOXA9 protein expression remained unaffected (Figure 4A). Moreover, a Cut&Run assay using CTCF antibody for chromatin immunoprecipitation confirmed loss of CTCF occupancy throughout the *HOXA9* locus, including CBS7/9 (Figure 4B). However, loss of CTCF occupancy did not correlate with a decrease in *HOXA7* or *HOXA9* expression at the mRNA level. Instead, long-term depletion of CTCF by auxin for 48 hours slightly increased the transcription of *HOXA7* and *HOXA9*. Upon washout of auxin from culture medium for an additional 48 hours, both *HOXA7* and *HOXA9* expression were restored to levels indistinguishable from those of the parental untreated cells (Figures 4C-4D). RNA-seq data collected from these three clones further confirmed the observation detected by Q-PCR (Figures S4B-S4C). Additionally, siRNA-mediated knock-down of CTCF in SEM cells did not change the transcription level of *HOXA7* or *HOXA9* (Figures 4E-4G). However, suppressing CTCF in human colorectal cancer cell line HCT116 notably reduced *HOXA7* and *HOXA9* expression (Figures S5A-S5C), consistent with the finding in MLLr AML cell line MOLM13 (24). Collectively, these data further confirmed the results of our CRISPR screening that CTCF is not a key regulator of *HOXA9* in MLLr B-ALL SEM and likely plays a role in regulating *HOXA9* transcription in a cell-type-specific manner.

**Figure 4.**
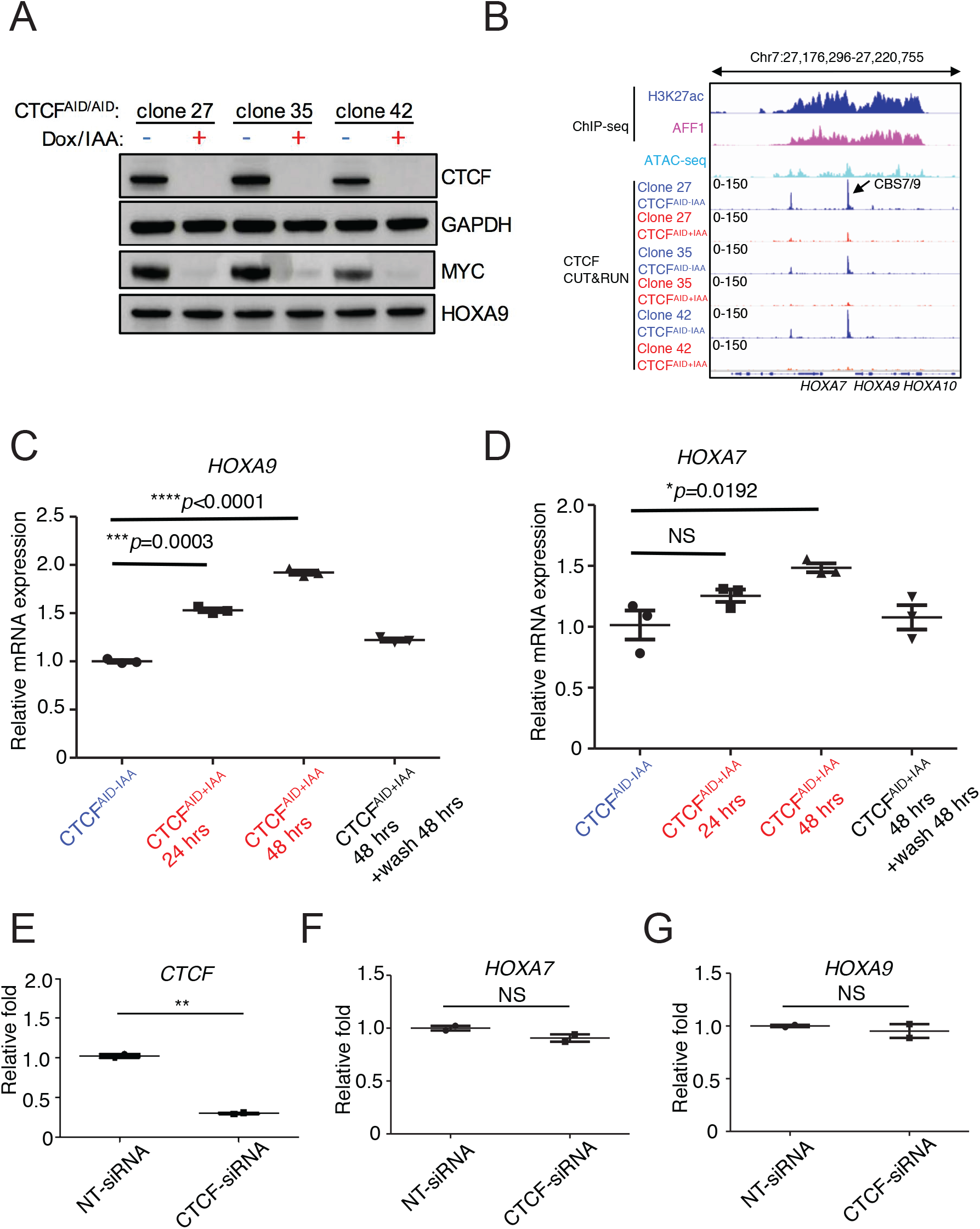
CTCF is dispensable for maintaining *HOXA9* expression in MLLr SEM cells. (A) Immunoblotting analysis of CTCF^AID^, MYC and HOXA9 in three bi-allelic knock-in clones 27, 35 and 42 with or without auxin (IAA) treatment. GAPDH was used as a loading control. (B) CTCF Cut&Run tracks shown at the selective viewpoint of the *HOXA9* locus where significant reduction of CTCF binding at CBS7/9 occurs following 48-hour IAA treatment in clones 27, 35, and 42. ChIP-seq tracks of CTCF, AFF1 and H3K27ac were included to indicate the open chromatin status of the locus. (C) Q-PCR analysis of *HOXA9* was conducted to monitor the transcriptional response to CTCF depletion for 24, 48 hours and washout from three biological replicates; clones 27, 35, and 42 (N=3). Data shown are means ± SEM from three independent experiments. *p < 0.05, **p < 0.01, ***p < 0.001, ****p < 0.0001, two-tailed Student’s *t* test. (D) Q-PCR analysis of *HOXA9* was conducted to monitor the transcriptional response to CTCF depletion for 24, 48 hours and washout from three biological replicates; clones 27, 35, and 42 (N=3). Data shown are means ± SEM from three independent experiments. *p < 0.05, two-tailed Student’s *t* test. (E) SEM cells were electroporated with CTCF-siRNA and NT-siRNA. Q-PCR was conducted 24 hours post electroporation to monitor *CTCF* expression. Data shown are means ± SEM from two independent experiments. **p < 0.01, two-tailed Student’s *t* test. (F) SEM cells were electroporated with CTCF-siRNA and NT-siRNA. Q-PCR analysis was conducted 24 hours post electroporation to monitor *HOXA7* expression. Data shown are means ± SEM from two independent experiments. *p < 0.05, two-tailed Student’s *t* test. (G) SEM cells were electroporated with CTCF-siRNA and NT-siRNA. Q-PCR was conducted 24 hours post electroporation to monitor *HOXA9* expression. Data shown are means ± SEM from two independent experiments. *p < 0.05, two-tailed Student’s *t* test.

### USF2 is required to maintain *HOXA9* expression in MLLr B-ALL

Aside from the positive controls confirmed from the CRISPR/Cas9 transcription factor screen in *HOXA9^P2A-mCherry^* cells, the top-ranked candidate among positive regulators was USF2. To further validate the CRISPR screen result and investigate the regulatory effect of USF2 on *HOXA9* expression, we individually delivered four lentiviral sgRNAs targeting *USF2* exons 1, 2, 7, and 9 into the *HOXA9^P2A-mCherry^* reporter line stably expressing Cas9. Similar to the results seen in sgENL targeted cells, *USF2* knock-down significantly decreased the mCherry fluorescence in a time-dependent manner compared to that of luciferase sgRNA-targeted control (sgLuc) (Figure 5A and Figure S6). Q-PCR and immunoblotting analysis further confirmed the concordant downregulation of both *HOXA9* and *mCherry* (Figures 5B-5C). Collectively, these data suggest that USF2 positively controls *HOXA9* expression in the MLLr cellular context. USF2 was reported to generally bind to a symmetrical DNA sequence (E-box motif) (5’CACGTG3’) in a variety of cellular promoters (47). Publicly available ChIP-seq data collected from human ES cells (48) suggested that USF2 can directly bind to the conserved E-box element at both *HOXA7* and *HOXA9* promoters (Figures S7A-S7B). A Cut&Run assay was performed in wildtype SEM cells to study genome-wide USF2 occupancy. Motif enrichment analysis by MEME-ChIP (49) revealed the top transcription factor binding motif is USF2 itself. Among the 36,299 binding peaks, about 25% contain a conserved USF2 binding motif (Figure 5D). Furthermore, the data confirmed the specific and strong binding of USF2 to an E-box element located in the *HOXA9* promoter in SEM cells (Figure 5E) suggesting USF2 could regulate *HOXA9* expression through interactions with its regulatory elements.

**Figure 5.**
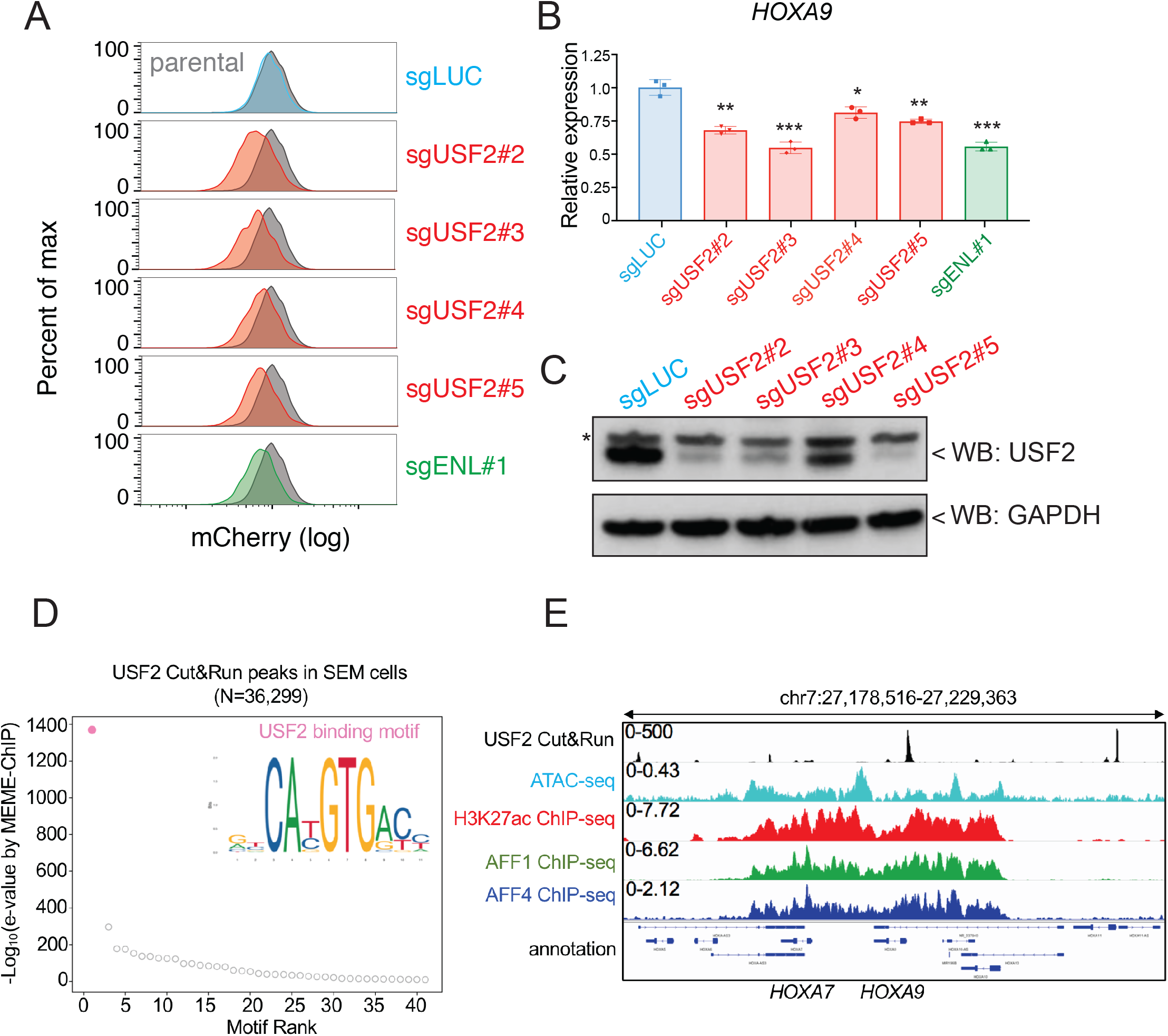
USF2 is required to maintain *HOXA9* expression in MLLr leukemia. (A) Flow cytometry analysis was performed at day 8 on the *HOXA9^P2A-mCherry^* cells targeted with lentiviral Cas9 and four sgRNAs against *USF2*. The sgENL targeted cells were used as positive controls while sgLuc targeted cells were used as negative controls. (B) Q-PCR analysis was conducted on the USF2 targeted cells to monitor the reduction of *HOXA9*. The sgENL targeted cells were used as positive controls while sgLuc targeted cells were used as negative controls. Data shown are means ± SEM from three independent experiments. *p < 0.05, **p < 0.01, ***p < 0.001, two-tailed Student’s *t* test. (C) Immunoblotting of USF2 in USF2 sgRNAs targeted cells. “*” denoted non-specific bands. (D) Motif analysis of genome-wide USF2-bound peaks by MEME-ChIP (49). (E) USF2 occupancy and ChIP-seq profiles of ATAC-seq, H3K27ac, AFF1 and AFF4 in *HOXA9* locus.

### USF2 is an essential gene in MLLr B-ALL by controlling *HOXA9* expression

To evaluate the importance of the USF2/HOXA9 axis in MLLr B-ALL progression, we sought to investigate the knockout phenotype of USF2 in MLLr B-ALL cells. A competition-based proliferation assay was performed by infecting SEM^Cas9^ cells with three individual lentiviral-mCherry-sgRNAs against USF2 (sgRNA-2, −3 and 5) at ~50% infection efficiency. The proportion of mCherry^+^ cells were monitored over a 12-day time course (days 3, 6, 9 and 12) to investigate the contribution of USF2 knock-down to cell survival. As a result, the proliferation-arrested phenotype was observed in all three sgRNA targeted cells but not in cells targeted with sgLuc (Figure 6A). Importantly, in SEM cells constitutively expressing ectopic retroviral mouse HOXA9 (SEM^HOXA9^) (Figure S8), USF2 knock-down had little effect on cell growth (Figure 6B), suggesting that HOXA9 is a functional and essential downstream gene of USF2 in regulating leukemia propagation. Next, to unbiasedly evaluate the survival dependency of USF2 in SEM cells, we conducted a dropout CRISPR/Cas9 screen by targeting 1,639 transcription factors. SEM cells infected with the pooled library of sgRNAs were collected at day 0 and day 12 to sequence for sgRNA distribution (Figure 6C). In accordance with prior genome-wide CRISPR screens and functional studies in B-ALL, many survival dependent genes were identified in the top 50 genes in our screen including *PAX5*, *DOT1L*, *ZFP64*, *YY1*, *MEF2C*, *MYC* and *KMT2A* (26, 29, 50, 51). USF2 was ranked as the top 24^th^ essential gene in MLLr SEM cells (Figure 6D). Taken together, these findings suggest that the USF2/HOXA9 axis plays a role in supporting MLLr B-ALL cell proliferation. In addition, a transcriptome analysis from the largest human B-ALL transcriptome cohort (N=1,988 patients) (26) identified *USF2* expression to be significantly correlated with *HOXA9* in MLLr-subtype patients (N=136 patients) (Figures 6E-6F and Figure S9A) highlighting that the USF2 and *HOXA9* regulation axis could have clinical relevance for patients in this specific subtype.

**Figure 6.**
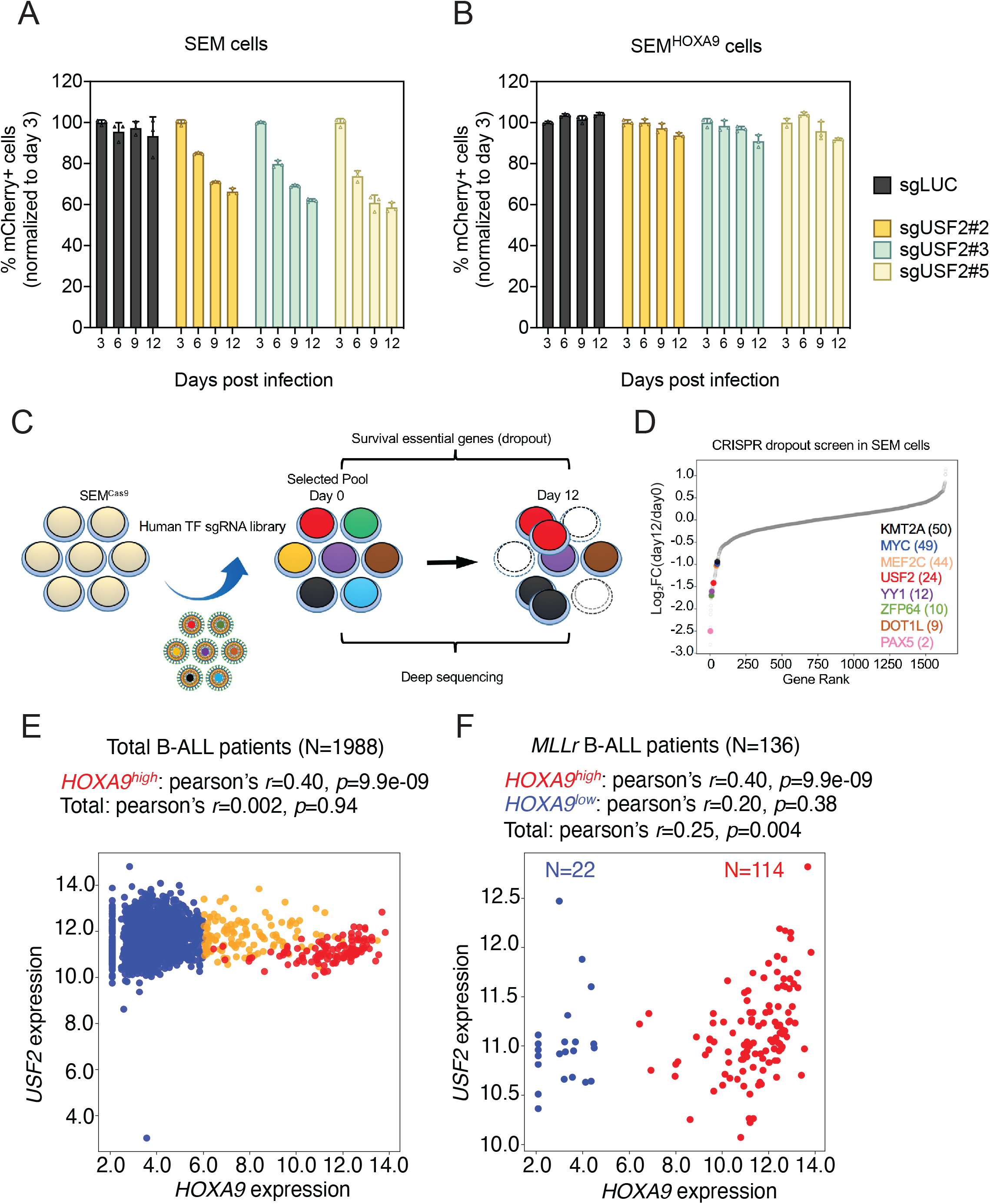
USF2 is a selectively essential gene in MLLr leukemia by controlling *HOXA9* expression. (A) Competitive proliferation assay was conducted by infecting SEM^Cas9^ cells with Lentiviral-mCherry-sgRNAs against luciferase (sgLuc) and USF2 (sgUSF 2-2, 2-3 and 2-5) at about 50% efficiency. The mCherry% was quantified every three days by flow cytometry to evaluate the growth disadvantage. The survival essential gene sgRPS19 was included as a positive control. (B) Rescued competitive proliferation assay was conducted by infecting SEM cells overexpressing ectopic HOXA9 with Lentiviral-mCherry-sgRNAs against luciferase (sgLuc) and USF2 (sgUSF2-2, 2-3 and 2-5) at about 50% efficiency. The mCherry% was quantified every three days by flow cytometry to evaluate the growth disadvantage. (C) Flow diagram of dropout CRISPR screening procedure. (D) Gene ranking of all transcription factors from dropout screening was illustrated. The enrichment score of seven sgRNAs against each transcription factor was combined by the MAGeCK algorithm. (E) Pearson’s correlation of transcriptional levels of *USF2* and *HOXA9* in a cohort of 1,988 B-ALL patients (26). (F) Pearson’s correlation of transcriptional levels of *USF2* and *HOXA9* in a cohort of 136 MLLr B-ALL subtype patients.

### USF1 and USF2 synthetically regulate *HOXA9* expression in MLLr leukemia

To further evaluate whether USF2 regulates HOXA9 expression in other MLLr leukemias, sgUSF2.2 was delivered into human AML cell line MOLM13 which carried MLL-AF9 translocation. Similar to those seen in SEM cells (Figures 7A and 7B), when USF2 protein was truncated by CRISPR targeting, HOXA9 expression was notably suppressed in MOLM13 cells (Figure 7C). Previously, other studies identified that a homolog protein USF1 shares the similar protein structure with USF2 (47, 52), recognizes the similarly conserved E-box elements across the genome. USF1 and USF2 are also able to form homo- or heterodimers (53–55), suggesting that these two proteins may function in synergy to regulate *HOXA9*. Interesting, in our HOXA9-reporter based CRISPR screen, USF1 was also among the top 50 positive regulator genes in our screen (49^th^) (Supplementary Table S2). Therefore, we co-delivered two sgRNAs against USF2 (sgUSF2.2) and USF1 (sgUSF1.3) to SEM *HOXA9^P2A-mCherry^* reporter line stably expressing Cas9. Notably, both of the flow cytometry analysis and Q-PCR confirmed that the *HOXA9* expression was suppressed to much lower level in double knockout of USF1 and USF2 compared with that in USF2 (Figures 7D and 7E). Collectively, USF1 and USF2 synthetically regulate *HOXA9* expression in human MLLr leukemia.

**Figure 7.**
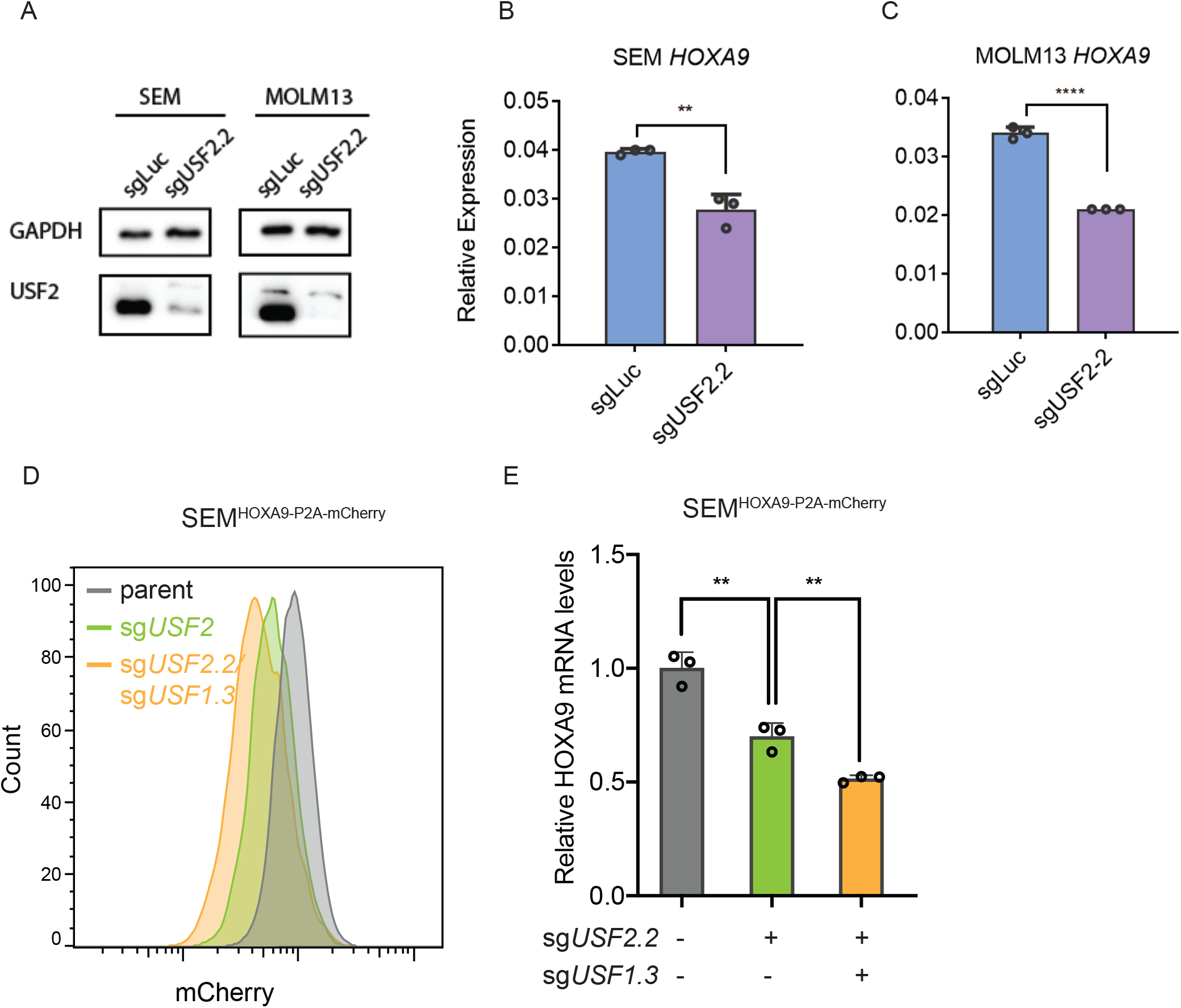
USF1 and USF2 synthetically regulate *HOXA9* expression in MLLr leukemia. (A) Immunoblotting analysis was conducted on the sgUSF2.2 targeted MOLM13 cells and control SEM cells to monitor the reduction of USF2 protein. (B) Q-PCR analysis was conducted on the sgUSF2.2 targeted MOLM13 cells and control SEM cells to monitor the reduction of USF2 protein. Data shown are means ± SEM from three independent experiments. **p < 0.01, ****p < 0.0001, two-tailed Student’s *t* test. (C) Flow cytometry analysis of mCherry was performed at day 9 on the *HOXA9^P2A-mCherry^* cells targeted with lentiviral Cas9 and sgRNAs against *USF1* (sgUSF1.3) and *USF2* (sgUSF2.2). (D) Q-PCR analysis was conducted on the USF1/USF2 targeted cells to monitor the reduction of *HOXA9*. Data shown are means ± SEM from three independent experiments. **p < 0.01, two-tailed Student’s *t* test.

### Non-coding regulation of *HOXA9* is associated with chromatin architecture in *HOXA* locus

In addition to protein-coding genes, noncoding DNA sequences also play important roles in regulating gene expression in *cis* (56–58). Our HOXA9 reporter system provides a robust platform for comprehensive profiling the functional *cis*-regulatory elements (CREs) that modulates HOXA9 transcription. To this end, we synthesized a 10,551-sgRNA array targeting the H3K27ac and ATAC-seq positive peaks defined in more than 500 human leukemia cell lines (CCLE) spanning the entire ~3 Mb region containing the *HOXA* cluster genes. An additional 100 non-targeting (NT) sgRNAs were also included as negative controls (Figure 7A). The lentiviral sgRNA library was transduced into *HOXA9^P2A-mCherry^* reporter line stably expressing dCas9-KRAB at a low multiplicity of infection (M.O.I<0.3), and then fractionated by flow cytometric sort for mCherry expression as described by human TF library screen previously (Figure 3). The mCherry^High^ and mCherry^Low^ populations in replicate screens were selected from the top or bottom 10% sorting gates and collected for deep sequencing to identify differentially represented sgRNAs, which indicated the corresponding targeted regions associated with transcriptional repression or activation of *p16^INK4A^*, respectively. Based on our observation that dCas9-KRAB and Cas9 effector both were efficient to identify functional *cis*-acting regulatory elements consistently (59), we sought to complement the dCas9-KRAB and sgRNA library screen by performing a parallel screen in *HOXA9^P2A-mCherry/+;Cas9^* stable SEM cells. Surprisingly, comparisons between dCas9-KRAB and wild-type Cas9 screens demonstrated poor correlation of global sgRNA distribution (*r*◻=◻0.011, *p*=0.23) (Figure 8B and 8C). At a stringent cut-off (an adjusted *p*-value of ≤0.01), we identified 26 differentially represented sgRNAs in the dCas9-KRAB screen (Figure 8D), mainly located on three hotspot regions including the positive control *HOXA9* promoter and adjacent intron 1, *HOXA10-AS* and *HOXA10* intron 1. Therefore, combining the *HOXA9^P2A-mCherry^* reporter cell line with CRISPR dCas9-KRAB screening identified two previously undiscovered regulatory elements in the *HOXA* locus (Figure S10C). In contrast, in Cas9-mediated non-coding screen, 524 sgRNAs designed to target the *HOXA6-10* region were significantly enriched in the mCherry^Low^ fraction, which was not observed in NT sgRNAs, *HOXA1-5* or *HOXA11-13* (Figure S10A). Notably, all of these enriched sgRNAs located at the hypomethylated valley identified by whole genome bisulfite sequencing (Figure 8D). Therefore, we hypothesize that Cas9-mediated double strand breaks at actively transcribed *HOXA6-10* locus may contribute to the transcription reduction of *HOXA9*. To test whether double strand breaks induced at other chromosomes would affect *HOXA9* transcription, we conducted the similar non-coding screen by infecting HOXA9P2A-mCherry; Cas9 cells with 2,049 sgRNAs targeting *CDKN2A*/*2B* locus. As a result, all of the sgRNAs were similarly distributed across mCherry^High^ and mCherry^Low^ populations (Figure S10B), indicating that the sorting-based screen did not bias the enrichment and the double strand breaks associated transcriptional regulation of *HOXA9* is locus specific. To further validate the screen result, five sgRNAs targeting CBS7/9, HOXA9 promoter, HOXA10-exon 1, HOXA9-intron 1 and HOXA10-AS and two control sgRNAs against Luciferase (LUC) and Rosa26 (ROSA) were individually infected into SEM^Cas9^ cells followed by antibiotic selection. As expected, Q-PCR confirmed the transcriptional repression of *HOXA9* in all five sgRNAs compared with negative control sgRNAs. Moreover, we also designed a sgRNA targeting on the *mCherry* coding cassette and delivered it into *HOXA9^P2A-^ mCherry;Cas9* reporter cell line and observed the specific transcriptional downregulation of *HOXA7* and *HOXA9*. Again, these data suggested that Cas9-mediated CRISPR targeting of actively transcribed hypomethylated *HOXA* cluster induced transcriptional repression of *HOXA9*.

**Figure 8.**
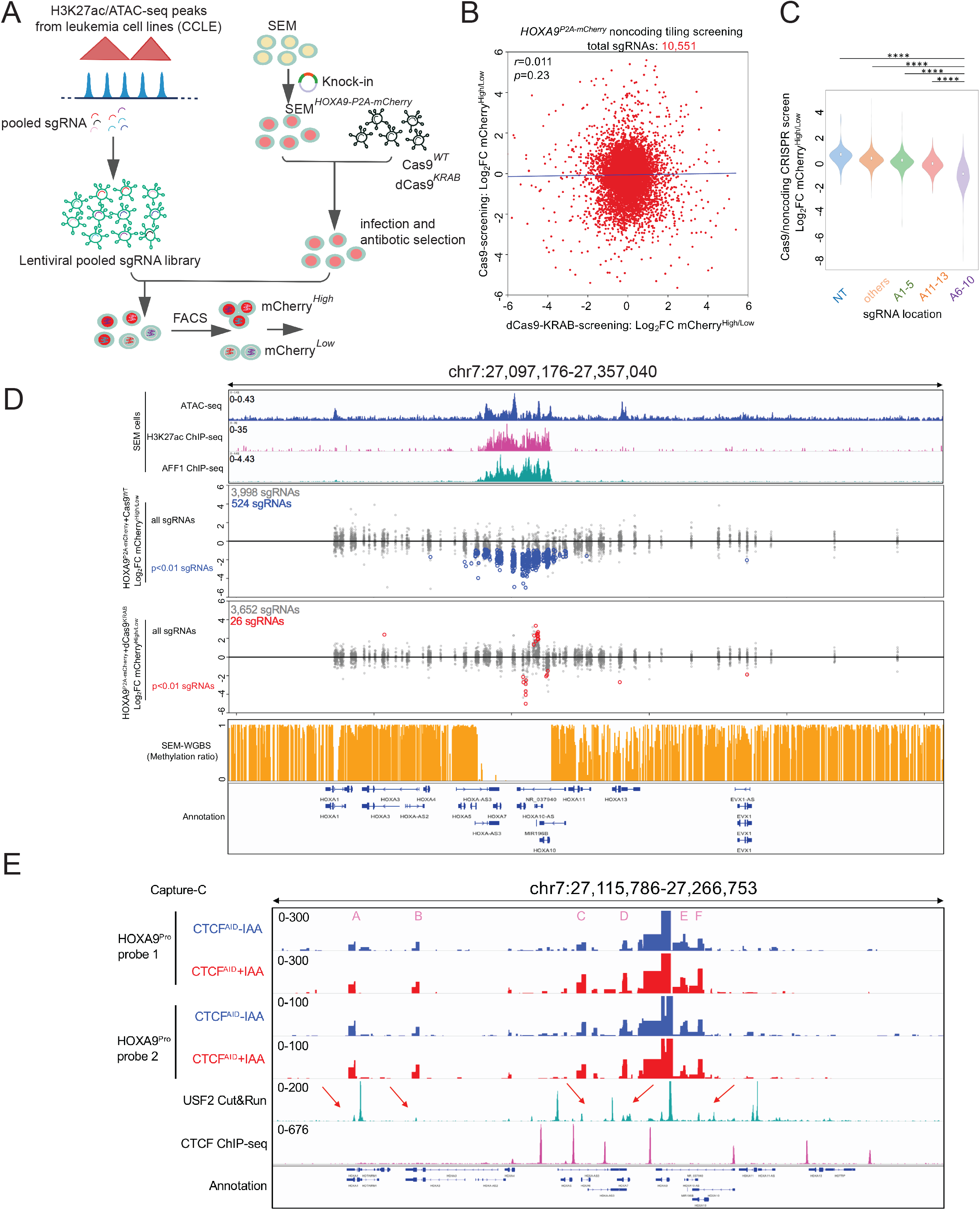
Non-coding regulation of *HOXA9* relies on chromatin architecture in *HOXA* locus. (A) Schematic diagram of dCas9-KRAB and Cas9 mediated non-coding screening in combination with *HOXA9^P2A-mCherry^* reporter allele. Total 10, 551 sgRNAs were designed and pooled in the non-coding library spanning the H3K27ac/ATAC-seq peaks identified from human leukemia cell lines (CCLE). (B) The global correlation of sgRNA distribution in dCas9-KRAB and Cas9 mediated screens. (C) Significant enrichment of sgRNAs targeting *HOXA6-10* regions was observed compared with other *HOXA* regions from the Cas9 mediated non-coding screen. (D) The global distribution of all sgRNAs in a selected region (chr7:27,097,176-27,357,040) of the *HOXA* locus from two screens in the *HOXA9^P2A-mCherry^* reporter cell line using dCas9-KRAB and Cas9. The sgRNAs with p-value cutoff of 0.01 were shown by blue in Cas9 and red in dCas9-KRAB mediated screens. Whole genome bisulfite sequencing indicated the hypomethylation status of *HOXA6-10* region in SEM cells. (E) Physical chromatin interactions between *HOXA9* and adjacent regions were detected by the next-generation Capture-C on parental SEM cells with or without CTCF protein. Two specific anchor probes (probe 1 and probe 2) were designed to hybridize to the *HOXA9* promoter, which identified six (A-F) interaction regions. The arrows indicate the Cut&Run peaks of USF2 in those HOXA9-interacted chromatin regions.

To test whether *HOXA9* promoter regulation requires long-distance chromatin interactions, we performed a high-resolution chromatin conformation capture assay, Capture-C, on a 3C library prepared from SEM cells with or without CTCF. Two biotinylated bait oligonucleotides were designed to hybridize to the *HOXA9* promoter. Strong enrichment of sequences at each bait site confirmed the efficiency of hybridization. In addition, six enriched regions (A-F) at *HOXA* cluster were identified as strong interacting regions, all of which overlap weak occupancy of USF2 shown by Cut&Run (Figure 8E). In consistent to our previous observation of CTCF acute depletion, chromatin interactions between *HOXA9* promoter and the six enriched regions was not affected upon CTCF degradation (Figure 8E). In summary, CTCF-independent high-order chromatin compaction likely plays an essential role in *HOXA9* regulation in MLLr B-ALL SEM cells.

## DISCUSSION

*HOX* genes are a cluster of genes strictly regulated in development by various transcription and epigenetic modulators. Mis-expression and dysregulation of *HOX* genes are frequently linked to human diseases, particularly cancer. Here, we focus on *HOXA9*, the aberrant expression of which is one of the most significant features in the most aggressive human leukemias. The *HOXA9^P2A-mCherry^* knock-in MLLr cell line derived in this study fully recapitulated transcriptional regulation of the endogenous gene. Previously, Godmin, *et al*. derived two mouse strains by delivering the in-frame GFP cassette to two different murine *Hox* genes, *Hoxa1* and *Hoxc13*, to visualize the proteins during mouse embryogenesis (60). Although this previous study certainly added to the repertoire of research tools available to investigate *HOXA*-related gene expression and gene function, our *HOXA9* reporter cell line provides a unique intrinsic cellular model with which to study transcriptional regulation of human *HOXA9* directly. Additionally, the CHASE-knock-in protocol developed to generate the *HOXA9* reporter is user-friendly, highly efficient, robust to reproduce and could be easily adapted to a wide variety of HOXA9-driven human leukemia cell models and other HOXA9-expressing cancer types.

In mammalian cells, each chromosome is hierarchically organized into hundreds of megabase-sized TADs (57, 61–63), each of which is insulated by the boundary elements. Within the TAD scaffold, promoter/enhancer physical contacts intricately regulate gene expression (64). Intra-TAD chromatin interactions can be facilitated by a pair of CTCF binding sites engaged in contact with each other when they are in a convergent linear orientation (65, 66). The *HOXA9* cluster is located on the TAD boundary, providing an opportunity to interact with neighboring genomic elements. However, because of the low resolution of publicly available Hi-C data and the lack of DpnI restriction enzyme sites within the *HOXA* gene cluster that are necessary to generate high-quality 3C libraries, the impact of chromatin interaction regulation of *HOXA9* remains unclear. Using a chromosome conformation capture-based PCR assay and CRISPR-mediated deletion of a minimal CTCF binding motif between *HOXA7* and *HOXA9* (CBS7/9), Luo and colleagues proposed that the CTCF boundary was crucial for higher-order chromatin organization by showing the depletion of CBS7/9 disrupted chromatin interactions and significantly reduced *HOXA9* transcription in MLLr AML MOLM13 cells with t(9;11) (24, 43). In our study, the loss-of-function results from auxin-inducible degradation of CTCF, siRNA-mediated CTCF knock-down, and the unbiased transcription factor screening suggested that CTCF is not required to maintain *HOXA9* expression in SEM cells with MLLr with t(4;11). We speculate that the discrepancy could be due to the following reasons. Although both cell lines carried the MLLr translocation as a driver oncogenic mutation, MOLM13 and SEM were classified as AML and B-ALL, respectively. Besides the lineage difference, SEM cells are also less sensitive to many well-known pharmaceutical inhibitors including JQ1 and DOT1L inhibitor. Therefore, we hypothesized that other as yet to be identified looping factors might be involved in the transcriptional regulation of the *HOXA9* locus in MLLr SEM cells, and that CTCF regulates HOXA9 expression in a cell-type-specific context.

By performing unbiased CRISPR screens designed to target 1,639 known human transcription factors in a *HOXA9^P2A-mCherry^* reporter cell line, we identified USF2 as a novel regulator of *HOXA9*. In addition, two known *HOXA9* regulators, *HOXA9* and *DOT1L* were identified among the top hits, supporting the reliable sensitivity of both the reporter system and the CRISPR screening strategy. USF2 is a ubiquitously expressed basic helix-loop-helix-leucine-zip transcription factor that generally recognizes E-box DNA motifs (47, 67, 68). USF1 and USF2 usually form homo- or heterodimers to modulate gene expression (53). Interestingly, USF1 was also enriched in our CRISPR screening. Moreover, the function of USF2 in controlling leukemia progression has not been reported. Our data from this study highlighted the plausible regulatory function of USF1/USF2 on *HOXA9* maintenance in MLLr B-ALL and AML cell lines, which would be an attractive target for therapeutic and mechanistic studies.

Our finding suggested that CTCF-independent high-order chromatin compaction likely contributed to this unique transcriptional regulation manner. We further revealed that candidate transcription factors identified from the CRISPR/Cas9 screen including USF2 and USF1, could also regulate *HOXA9*, thereby providing a more comprehensive understanding about how the *HOXA9* locus is regulated in human cancer cells. Further mechanism studies will be exploited in the future. Given the well-recognized role of *HOXA9* in hematopoietic malignancy, we anticipate the *HOXA9* reporter cells will advance many lines of investigation, including drug screening and the identification of concordant epigenetic modifiers/transcription factors that are required for activation and maintenance of *HOXA9* expression in leukemia progression. Collectively, these efforts would clarify the molecular mechanisms underlying aberrant *HOXA9* activation in leukemias, thus providing the foundation to develop clinically relevant therapies to target the expression and/or function of *HOXA9* in leukemia patients.

## METHODS and MATERIALS

### Cell culture

SEM cells (ACC-546, DSMZ) and MOLM13 (ACC-554, DSMZ) were maintained in RPMI-1640 medium (Lonza) containing 10% fetal bovine serum (FBS) (HyClone), and 1% penicillin/streptomycin (Thermo Fisher Scientific) at 37°C, 5% CO_2_ atmosphere and 95% humidity. Basal medium for culturing 293T cells is DMEM (HyClone). All passages of cells used in this study were mycoplasma-free. Cell identity was confirmed by STR analysis.

### Vector construction

A pair of oligomers containing a 20-bp sgRNA (5’-AAAGACGAGTGATGCCATTT-3’) sequence targeting the surrounding genomic segment of *HOXA9* stop codon was synthesized (Thermo Fisher Scientific) and cloned into the all-in-one vector, pSpCas9(BB)-2A-GFP (Addgene #48138) between *BsmBI* sites. Correct clones were screened and confirmed by Sanger sequencing with the U6-Forward sequencing primer (5’-GAGGGCCTATTTCCCATGAT-3’). To construct a CHASE-knock-in donor vector delivering a *P2A-mCherry* DNA segment to the endogenous *HOXA9* locus, a two-step cloning protocol was used. The ~800-bp 5′ and 3’ homology arm (HA) flanking the endogenous sgRNA target was amplified from SEM cells. The 5’ HA PCR primer sequences are 5’-GGCCGATTCCTTCCACTTCT-3’ and 5’-TCACTCGTCTTTTGCTCGGT-3’, and the 3’ HA PCR primer sequences are 5’-ACCGAGCAAAAGACGAGTGA-3’ and 5’-CACTGTTCGTCTGGTGCAAA-3’. The *P2A-mCherry* DNA fragment was amplified from p16^INK4A^-P2A-mCherry knock-in donor vector (29) using a pair of primers containing overlapping sequences of 5’ HA or 3’ HA for in-fusion cloning (forward primer: 5’-AAGACCGAGCAAAAGACGAGGGATCCGGCGCAACAAACTT-3’; reverse primer: 5’-AATAAGCCCAAATGGCATCACTTGTACAGCTCGTCCATGC-3‘). The 5’ HA-P2A-mCherry-3’ HA in-fusion cloning product was further supplemented with 23-bp target sgRNA and PAM sequences at both 5’ and 3’ ends through PCR amplification using primers 5’-AAAGACGAGTGATGCCATTTGGGATGAGGCTGCGGGCGAC-3’ and 5’-AAAGACGAGTGATGCCATTTGGGTATATATACAATAGACAAGACAGGAC-3’. The cloning PCR reactions were performed using Q5 High-Fidelity DNA Polymerase (New England Biolabs # M0491L), and the cycling parameters were as follows for all cloning: 98°C for 30 s, followed by 98°C for 15 s, 72°C for 20 s, and 72°C for 30 s per kb for 40 cycles. The final PCR product was conducted into TOPO cloning vector (Thermo Fisher Scientific #450641). Sanger sequencing was performed to ensure that the knock-in DNA was cloned in-frame with the HAs. The Lenti-Cas9-Blast plasmid (#83480) and the Lenti-Guide-Puro plasmid (#52963) were purchased from Addgene. For candidate validation of CRISPR screen, sgRNA sequences against *DOT1L*(5’-TCAGCTTCGAGAGCATGCAG-3’), *ENL*(5’-TCACCTGGACGGTGCACTGG-3’), *USF2* (#2: 5’-AGAAGAGCCCAGCACAACGA-3’, #3: 5’-TGTTTTCCGCAGTGGAGCGG-3’, #4: 5’-CCGGGGATCTTACCTGGCGG-3’, and #5: 5’-CAGCCACGACAAGGGACCCG-3’) were cloned into an in-house-made Lenti-Guide-Puro-IRES-CFP vector. The sgRNA sequence against *USF1* (3#, 5’-CTATACTTACTTCCCCAGCA-3’) was cloned into an in-house-made LRNeo-2.1 vector in which the mCherry-expressing cassette of LRCherry2.1 (Addgene #108099) was replaced by Neomycin. For competitive proliferation assay, sgRNAs against Luciferase (Luc)(5’-CCCGGCGCCATTCTATCCGC-3’) and USF2 (#2, #3 and #5 as above) were cloned into mCherry-expressing LRCherry2.1 (Addgene #108099) vector.

### Generation of a *HOXA9^P2A-mCherry^* reporter allele

SEM were electroporated by using the Nucleofector-2b device (Lonza) with the V-kit and program X-001. For *HOXA9^P2A-mCherry^* knock-in delivery, 2.5 μg of the donor plasmid and 2.5 μg of the CRISPR/Cas9-HOXA9-C-terminus-sgRNA all-in-one plasmid were used for 5 million SEM cells. Twenty-four hours after transfection, cells were sorted for the GFP fluorescent marker linked to Cas9 expression vector to enrich the transfected cell population. After the sorted cells recovered in culture for up to 3 weeks, a second sort was performed to select cells for successful knock-in by sorting for cells expressing the knock-in mCherry fluorescent marker. Two weeks later, a third sort was repeated based on the selection mCherry expressing cells.

### Characterization of successful knock-in events by PCR and Sanger Sequencing

DNA from single-cell-derived bacterial or cell colonies was extracted with a Quick-DNA Miniprep Kit (Zymo #D3025). Combinatorial primer sets designed to recognize the 5′ and 3′ knock-in boundaries were used with the following PCR cycling conditions: 98°C for 2 mins, followed by 40 cycles of 98°C for 30 s and 68°C for 60 s. The sequences for genotyping primers are provided in Supplemental Table 1. After electrophoresis, the bands that were at the expected size were cut out, purified, and sequenced with two specific primers (Supplementary Table S1).

### CRISPR library construction and screening

A set of ~10,000-sgRNA oligos that target 1,639 human transcription factors were designed for array-based oligonucleotide synthesis (CustomArray). Unique binding of each sgRNA was verified by sequence blast against the whole human genome. In the sgRNA pooled library, seven gRNAs against each of the 1,639 human transcription factors were obtained from validated sgRNA libraries published previously (69–77). The synthesized oligo pool was amplified by PCR and cloned into LentiGuide-Puro backbone (#52963) by in-fusion assembly (Clontech #638909). The *HOXA9^P2A-mCherry^* reporter cell line was overexpressed with lentiviral Cas9 followed by infection of pooled sgRNA library at low M.O.I (~0.3). Infected cells were selected by blasticidine and puromycin and later sorted for mCherry^High^ and mCherry^Low^ populations between days 10-12. The sgRNA sequences were recovered by genomic PCR analysis and deep sequencing using MiSeq for single-end 150-bp read length (Illumina). The primer sequences used for cloning and sequencing are listed in Supplementary Table S1. The sgRNA sequences are described in Supplementary Table S2. High-titer lentivirus stocks were generated in 293T cells as previously described (78).

### Data analysis of CRISPR screening

The raw FASTQ data were de-barcoded and mapped to the original reference sgRNA library. The differentially enriched sgRNAs were defined by comparing normalized counts between sorted cells in the top 10% and those in the bottom 10% of mCherry-expressing bulk populations. Two independent replicate screenings were performed with the *HOXA9^P2A-mCherry^* reporter cell line stably expressing Cas9. Normalized counts for each sgRNA were extracted and used to identify differentially enriched sgRNA by DESeq2 (42). The combined analysis of seven sgRNAs against each human transcription factor was conducted by using the MAGeCK algorithm (41). Detailed screening results were included in Supplementary Table S2.

### Fluorescence imaging and analysis

0.1% of DMSO (vehicle control) or 10 doses of SGC0946 with a half log scale (0.3 nM-10 μM) were first dispensed into 384-well plates (in quadruplicate, 4 wells per dose). Suspension-cultured SEM cells were immediately plated into the 384-well plate (20,000 cells / well). Six days after drug treatment, the cells were fixed with 4% paraformaldehyde for 10 mins at room temperature, followed by Hoechst staining for 15 mins at room temperature. Fluorescence images (Hoechst and mCherry) were taken by a CellVoyager 8000 high content imager (Yokogawa). The acquired images were processed by using the Columbus Image Data Storage and Analysis system (Perkin Elmer) to count the number of positive cells and measure fluorescent intensity. To determine the changes of mCherry intensity in SEM expressing *HOXA9^P2A-mCherry^*, we measured average mCherry intensity of four fields per well and normalized to vehicle (0.1% DMSO) treated control. Wild-type SEMs with no fluorescence were included as negative controls.

### Cut&Run assay

Cut&Run assay was conducted following the protocol described previously (79). In brief, three million cells were collected for each sample. The USF2 antibody (NBP1-92649, Novus) was used at a 1:100 dilution for immunoprecipitation. Library construction was performed using the NEBNext UltraII DNA Library Prep Kit from NEB (E7645S). Indexed samples were run using the Illumina Next-seq 300-cycle kit. Cut&Run raw reads were mapped to genome hg19. by bowtie 2.3.4 with default parameter. The mapping file were converse to .bw file by bamCoverage (80, 81).

### Flow cytometry

Suspension-cultured SEM were collected by centrifugation at 800X*g*, filtered through a 70-μm filter, and analyzed for mCherry on a BD FACS Aria III flow cytometer with a negative control. The 4,6-diamidino-2-phenylindole (DAPI) staining was conducted prior to sorting to exclude dead cells.

### Inhibitor treatment

SEM cells were seeded at a density of 1×10^5^ cells/mL in medium supplemented with DMSO vehicle or different doses (from 0.5 μM to 15 μM) of the DOT1L inhibitor SGC0946 (MedChemExpress #HY-15650). Medium was replaced every three days, and fresh inhibitor was added. At day-6 post treatment, cells were collected for flow cytometry analysis and RNA extraction.

### Fluorescence *in situ* hybridization

An ~800-bp purified *P2A-mCherry* DNA fragment was labeled with a red-dUTP (AF594, Molecular Probes) by nick translation, and a *HOXA9* BAC clone (CH17-412I12/7p15.2) was labeled with a green-dUTP (AF488, Molecular Probes). Both of labeled probes were combined with sheared human DNA and independently hybridized to fix the interphase and metaphase nuclei derived from each sample by using routine cytogenetic methods in a solution containing 50% formamide, 10% dextran sulfate, and 2XSSC. The cells were then stained with DAPI and analyzed.

### Quantitative real-time PCR

Total RNA was collected by using TRIzol (Thermo Fisher Scientific #15596026) or Direct-zol RNA Miniprep Kit (Zymo #R2052). Reverse transcription was performed by using a High-Capacity cDNA Reverse Transcriptase Kit (Applied Biosystems #4374966). Real-time PCR was performed by using FAST SYBR Green Master Mix (Applied Biosystems #4385612) in accordance with the manufacturer’s instructions. Relative gene expression was determined by using the ΔΔ-CT method (82). All Q-PCR primers used in this study are listed in Supplementary Table S1.

### Competitive proliferation assay

For evaluating the impact of USF2 sgRNAs on leukemia expansion, cell cultures were lentivirally transduced with individual USF2 sgRNAs in mCherry expressing vector, followed by measurement of the mCherry-positive percentage at various days post-infection using flow cytometry. The rate of mCherry-positive percentage was normalized to that of Day 3 and declined over time, which was used to infer a defect in cell accumulation conferred by a given sgRNA targeting USF2 relative to the uninfected cells in the same culture.

### Statistics

All values are shown as the mean[±[SEM. Statistical analyses were performed with GraphPad Prism software, version 6.0. *P*-values were calculated by performing a two-tailed *t*-test.

## ACKNOWLEDGMENTS

We gratefully acknowledge the staffs of the Hartwell Sequencing, Cytogenetics, Flow Cytometry and Cell Sorting Shared Resource facility within the Comprehensive Cancer Center of St. Jude Children’s Research Hospital. We thank Li and Lu laboratory members for critical comments and discussion. We thank Dr. Cherise Guess for helping with scientific editing.

## FUNDING

This work was funded in part by the St. Jude Comprehensive Cancer Center development fund NCI-5P30CA021765-37 from the National Cancer Institute (C.L.), the American Lebanese Syrian Associated Charities (C.L), startup funds from the Division of Hematology/Oncology at the University of Alabama at Birmingham (R.L.), Leukemia Research Foundation (R.L.), Mary Ann Harvard Award from the Young Supporters Board of the O’Neal Comprehensive Cancer Center (R.L.), and R35-GM118041 from the National Institute of General Medical Sciences (T.C.).

## AUTHOR CONTRIBUTIONS

Conceptualization: C.L. and R.L.; Methodology: H.Z., Y.Z., J.H., S.W., L.Z., J.A., Y.S., Y.Z., T.C., H.L., B. Xu., R.L. and C.L.; Investigation: H.Z., Y.Z., J.H., S.W., L.Z., J.A., Y.Z., R.L. and C.L.; Software and formal analysis: Y.Z.; Writing, Review, and Editing: R.L. and C.L.; Supervision, project administration, and funding acquisition: R.L. and C.L.

## DECLARATION OF INTERESTS

The authors declare no competing interests

## AVAILABILITY OF DATA AND MATERIALS

All plasmids created in this study will be deposited to Addgene. Raw data collected from Cut&Run were deposited at NCBI GEO (GSE140664). Raw data collected from CRISPR screening were included in supplementary Table S2. Publicly available dataset used in this study were cited accordingly. Figures 1E and S4B: GSE120781; Figure 4B: GSE126619; Figures 6E and 6F: European Genome-phenome Archive (EGA) under accession number EGAS00001003266, EGAS00001000654, EGAS00001001952, EGAS00001001923, EGAS00001002217 and EGAS00001000447; Figure S1: GSE13159.

## ETHICS APPROVAL AND CONSENT TO PARTICIPATE

Not applicable

## CONSENT FOR PUBLICATION

Not applicable

## SUPPLEMENTAL FIGURE LEGENDS

**Figure S1.**
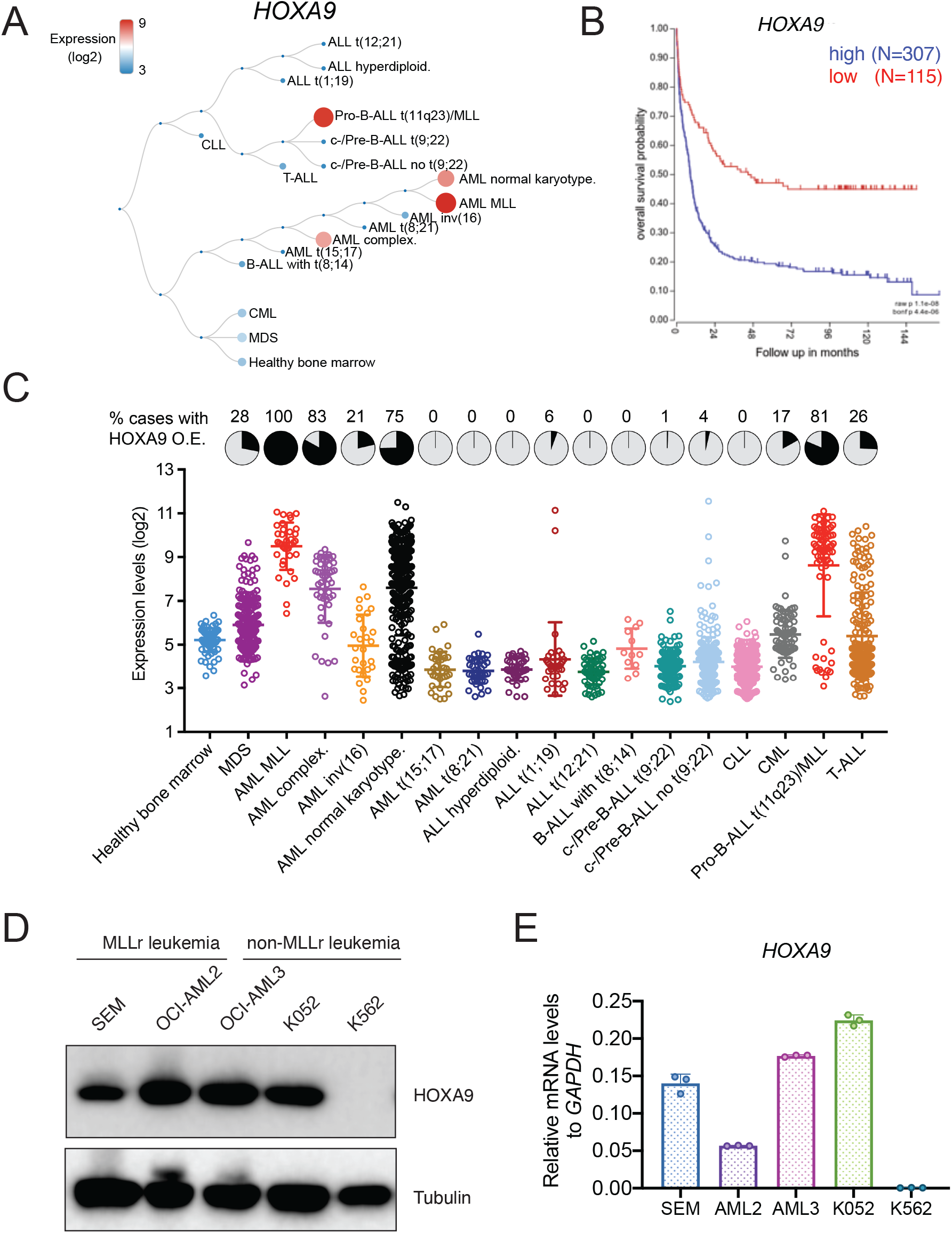
*HOXA9* expression profiling in leukemia. (A) *HOXA9* expression in different leukemia lineages (GSE13159). (B) Kaplan-Meier survival curve indicated the poor outcome associated with high HOXA9 expression (GSE13159). (C) *HOXA9* expression was revealed by leukemia subtypes in MILE leukemia study cohort (bloodspot). (D) HOXA9 protein level was assessed by immunoblotting in MLLr and non-MLLr leukemia cell lines. (E) *HOXA9* mRNA level was assessed by Q-PCR in MLLr and non-MLLr leukemia cell lines.

**Figure S2.**
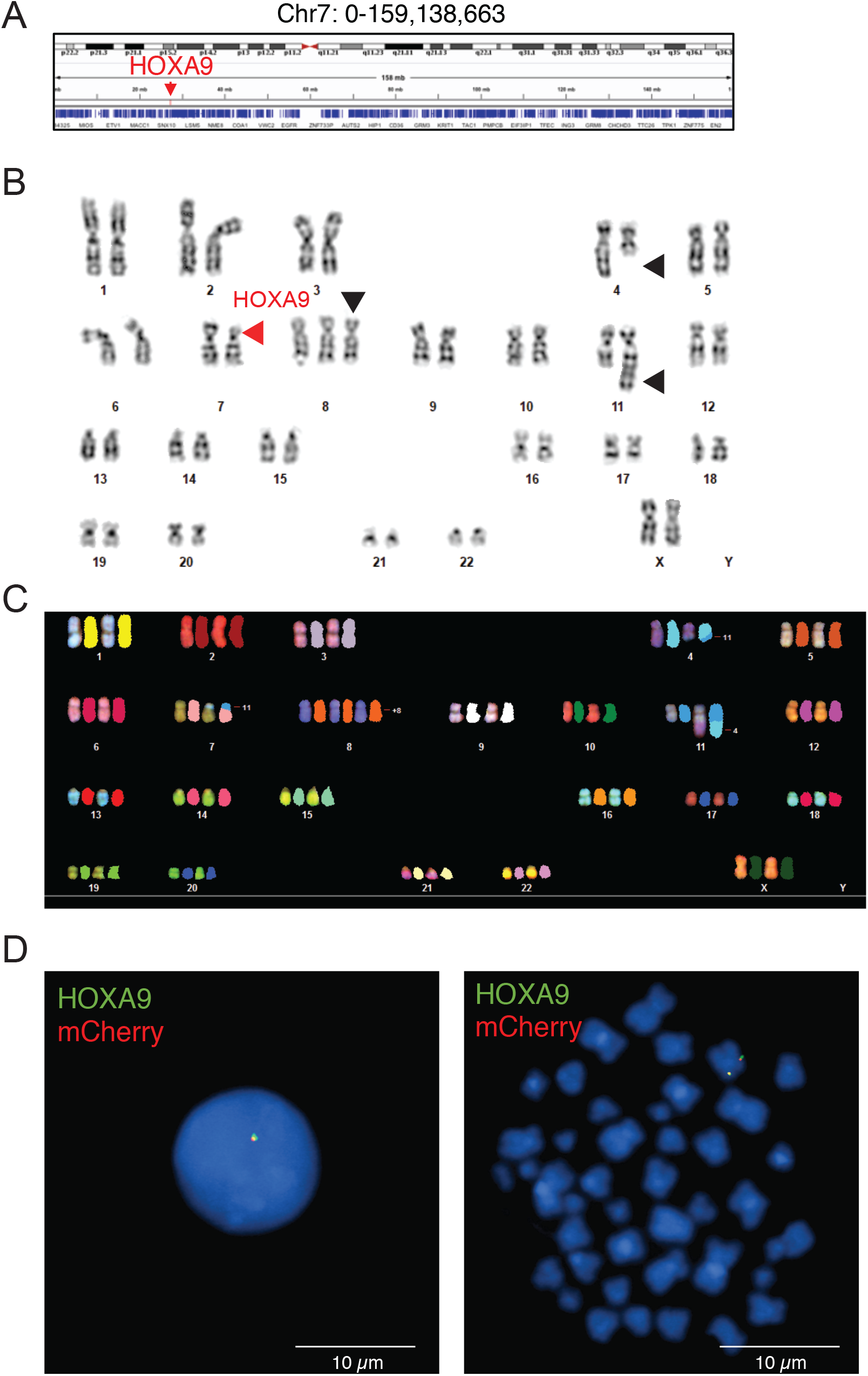
Cytogenetic characterization *HOXA9* knock-in allele in MLLr SEM cells. (A) The genomic *HOXA9* location was highlighted in human chromosome 7. (B) Karyotype analysis of parental MLLr SEM cells indicating the mono-allelic deletion of partial segment in chromosome 7 containing the *HOXA* cluster (red arrow). Black arrows indicated other chromosome alterations including t4,11 translocation and trisomy 8. (C) Chromosome analysis of spectral karyotyping (SKY) was conducted by using a commercially prepared SKY probe from Applied Spectral Imaging (Carlsbad, CA) on HOXA9 reporter cells. Translocation between chr4 and chr11, trisomy 8 and micro-deletion of chr7 was confirmed. (D) FISH analysis confirming the co-localization of *HOXA9* and mCherry in targeted cells at interphase (left) and metaphase (right). The *P2A-mCherry* DNA was labeled with a red-dUTP by nick translation, and an *HOXA9 BAC* clone was labeled with a green-dUTP. The cells were then stained with 4,6-diamidino-2-phenylindole (DAPI) to visualize the nuclei. A representative cell image is shown for the pattern of hybridization (pairing of red and green signals).

**Figure S3.**
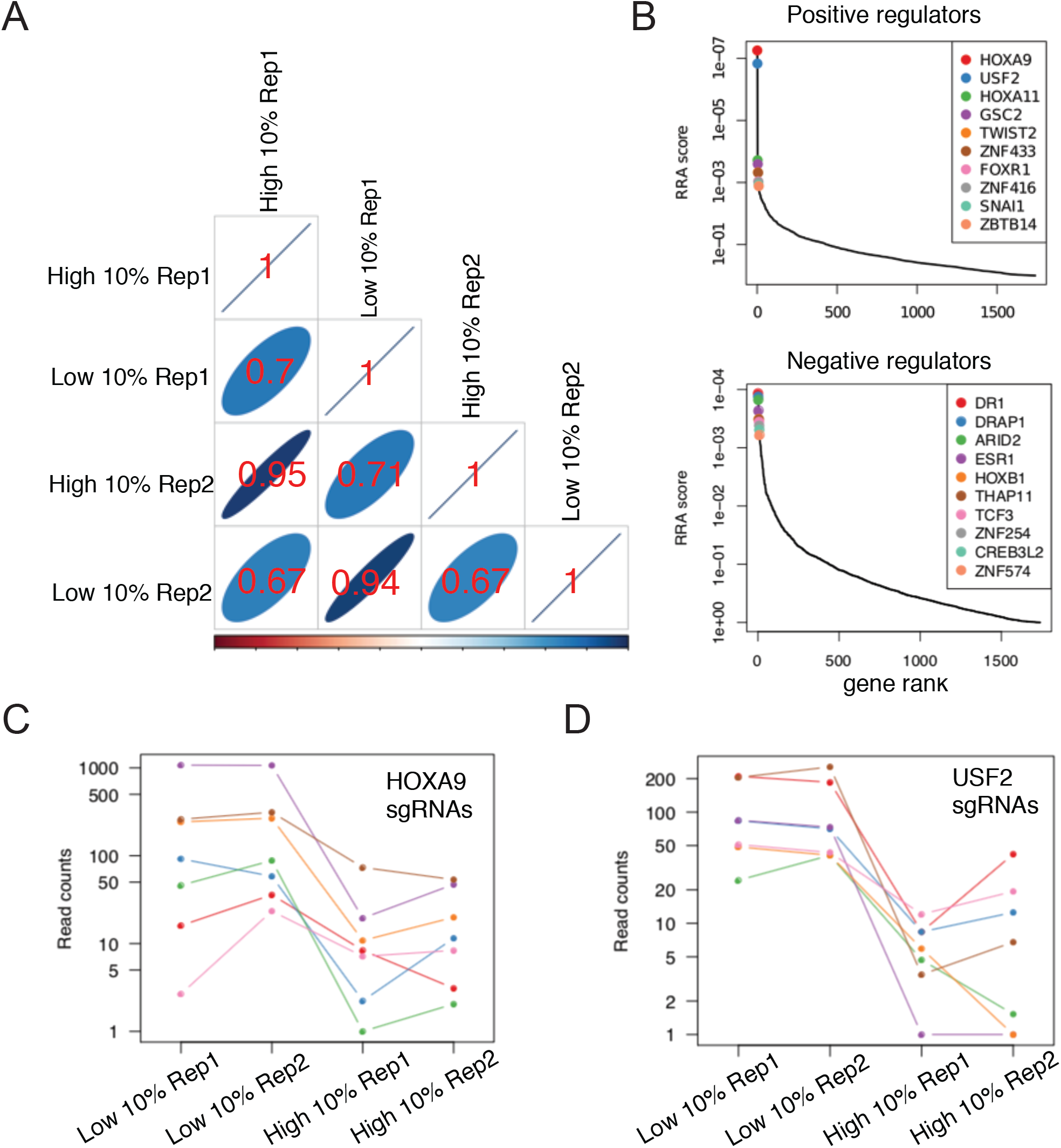
Cytogenetic characterization of the *HOXA9* knock-in allele in MLLr SEM cells. (A) Pearson’s correlation of normalized sgRNA counts in mCherry^High^ and mCherry^Low^ sorted populations. (B) Gene ranking of the top 10 positive and negative candidate regulators of *HOXA9* enriched from screening analysis by MAGeCK algorithm. (C) Normalized sgRNA count distribution of each of seven sgRNAs against *HOXA9*. (D) Normalized sgRNA count distribution of each of seven sgRNAs against *USF2*.

**Figure S4.**
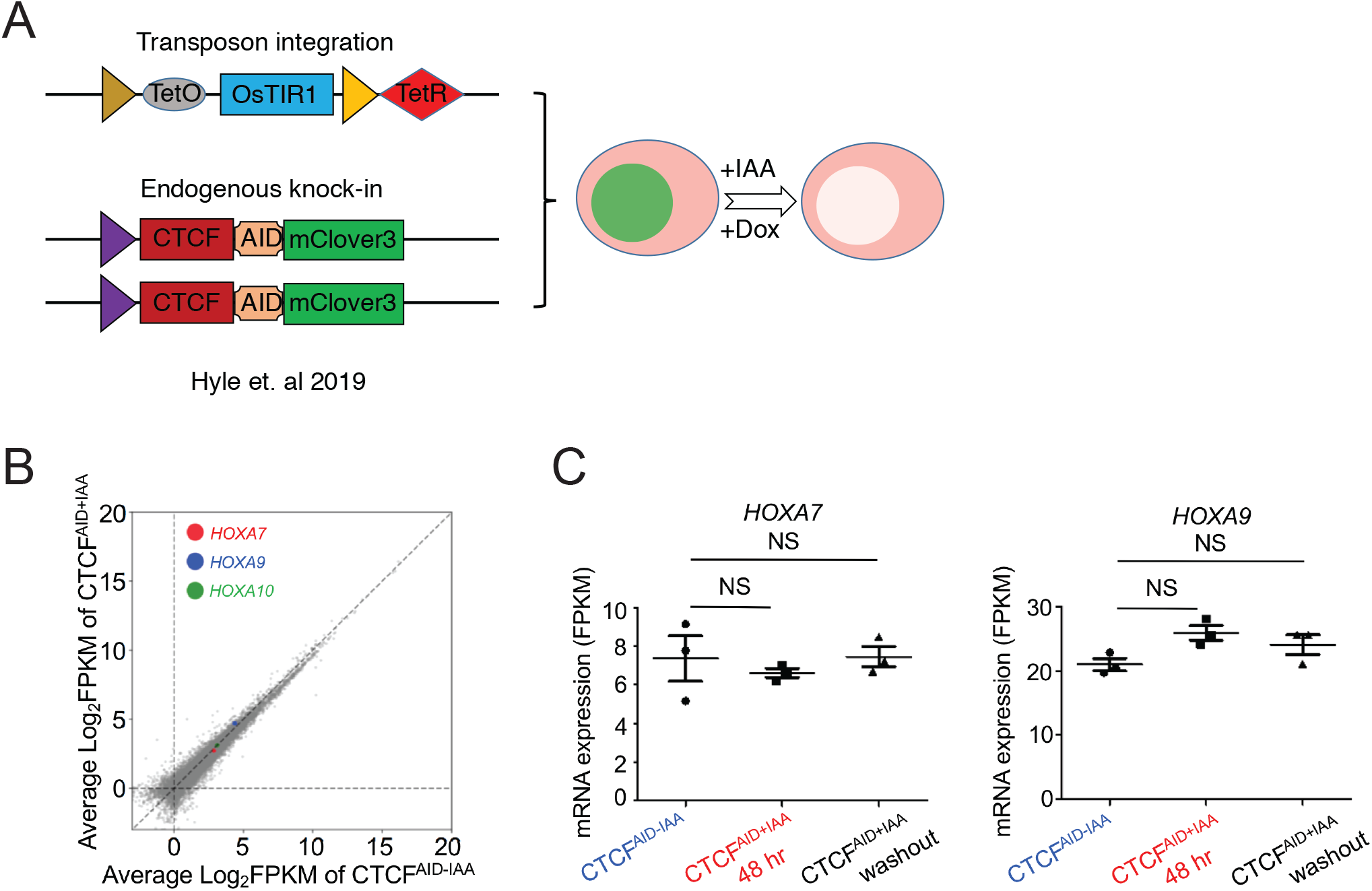
Auxin-inducible degradation of CTCF does not affect *HOXA9* expression in SEM cells. (A) Flow diagram of auxin-inducible degradation model to acutely deplete endogenous CTCF protein. Dox, doxycycline; IAA: auxin. (B) RNA-seq profiles of *HOXA7*, *HOXA9* and *HOXA10* in CTCF depleted SEM cells. (C) Quantification of *HOXA7*, *HOXA9* and *HOXA10* levels in three knockin clones of CTCF depleted SEM cells.

**Figure S5.**
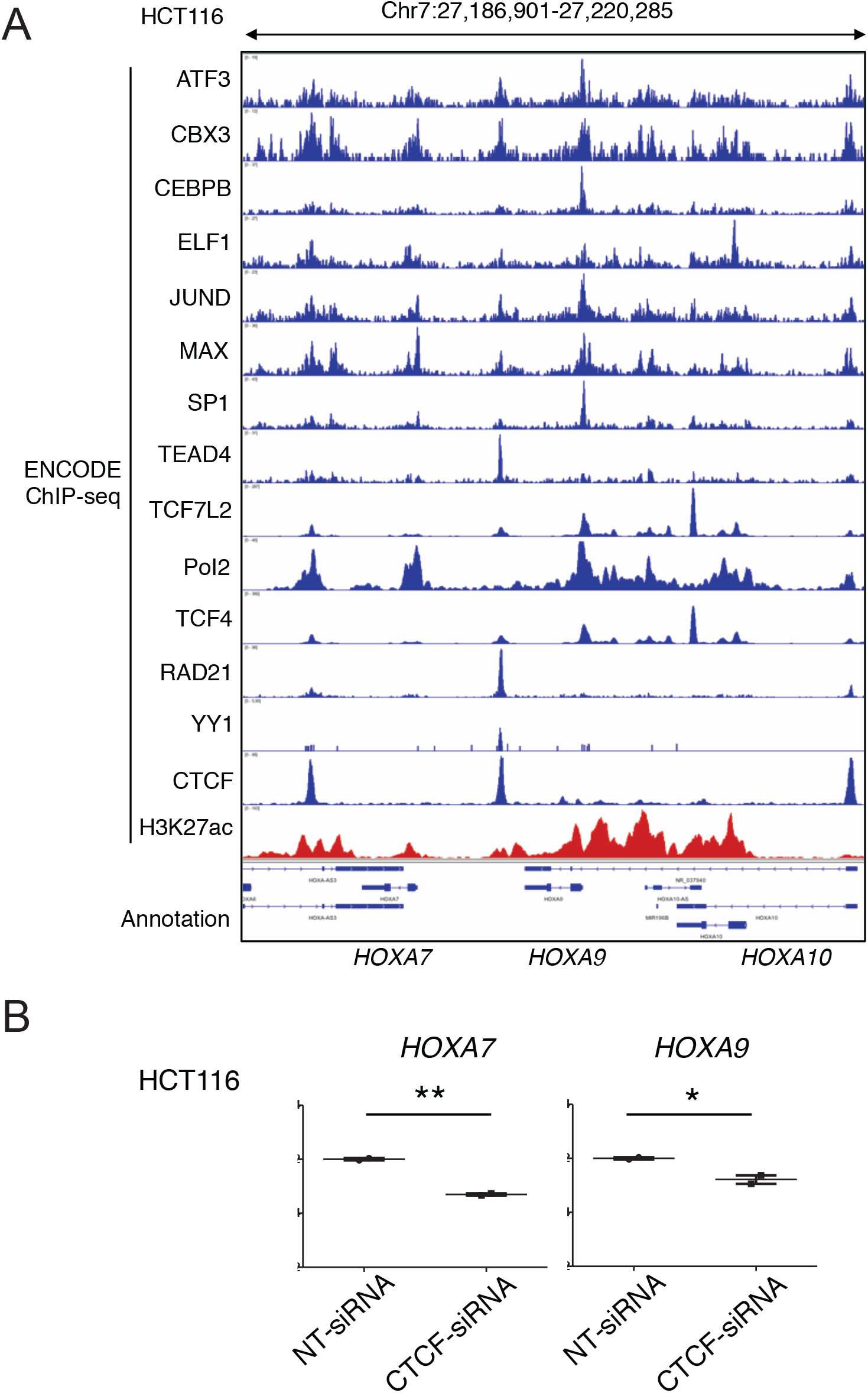
CTCF regulates *HOXA9* expression in human colorectal cancer HCT116 cells. (A) ChIP-seq tracks from publicly available ENCODE dataset demonstrated the enriched transcription factor occupancy at CBS7/9 in HCT116 cells RNA-seq profiles of *HOXA7*, *HOXA9* and *HOXA10* in CTCF depleted SEM cells. (B) Q-PCR analysis of HOXA7 and HOXA9 in HCT116 cells transfected with CTCF-siRNAs and NT-siRNAs for 48 hours. Data are means ± SEM from two independent experiments. *p < 0.05, **p < 0.01, Student’s *t* test.

**Figure S6.**
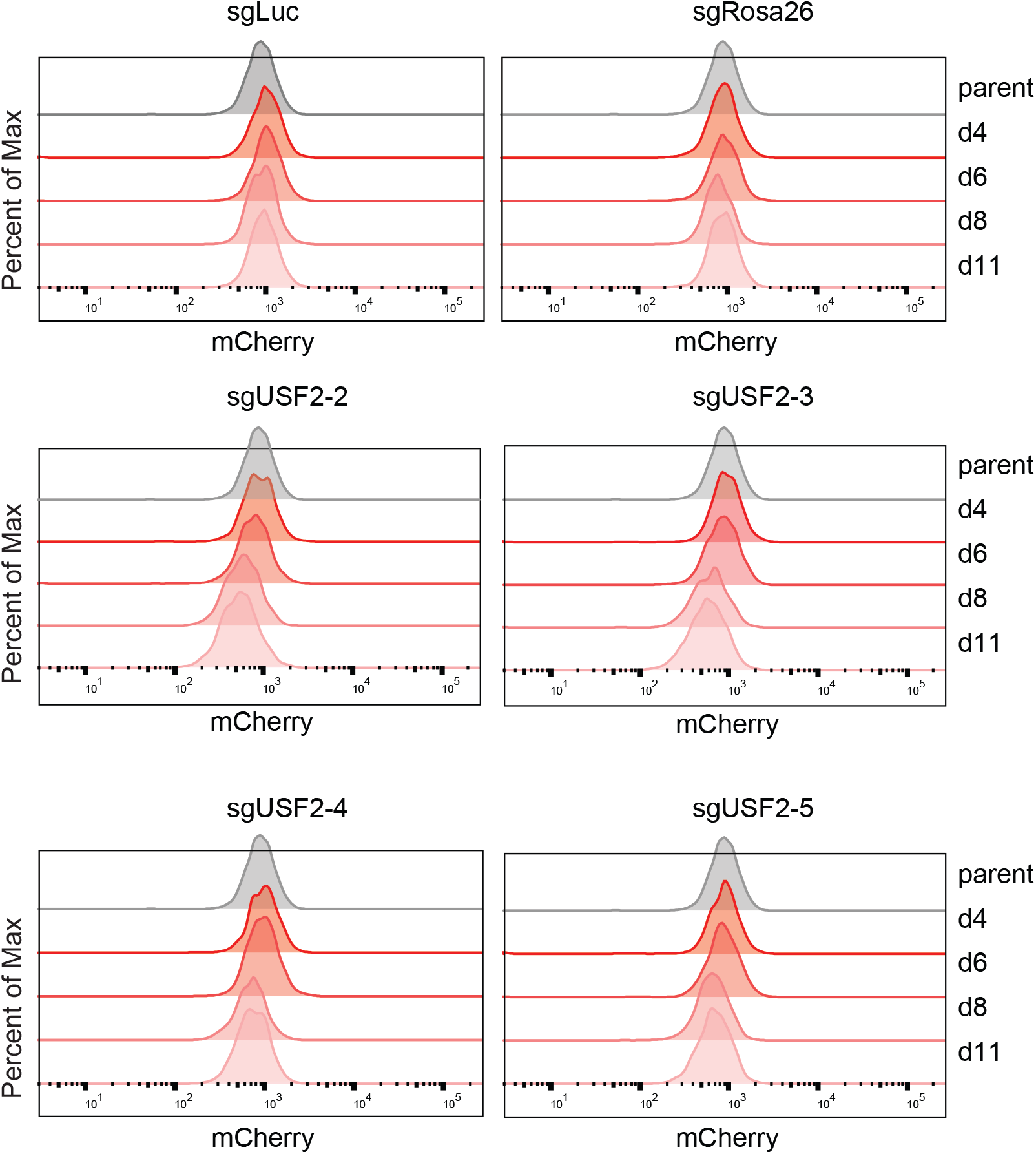
Time-course knocking down of *USF2* and consequent *HOXA9* expression analysis. Flow cytometry analysis was performed at day 0, 4, 6, 8 and 11 on the *HOXA9^P2A-mCherry^* cells targeted with lentiviral Cas9 and four sgRNAs against *USF2*. The sgLuc and sgRosa26 targeted cells were included as negative controls.

**Figure S7.**
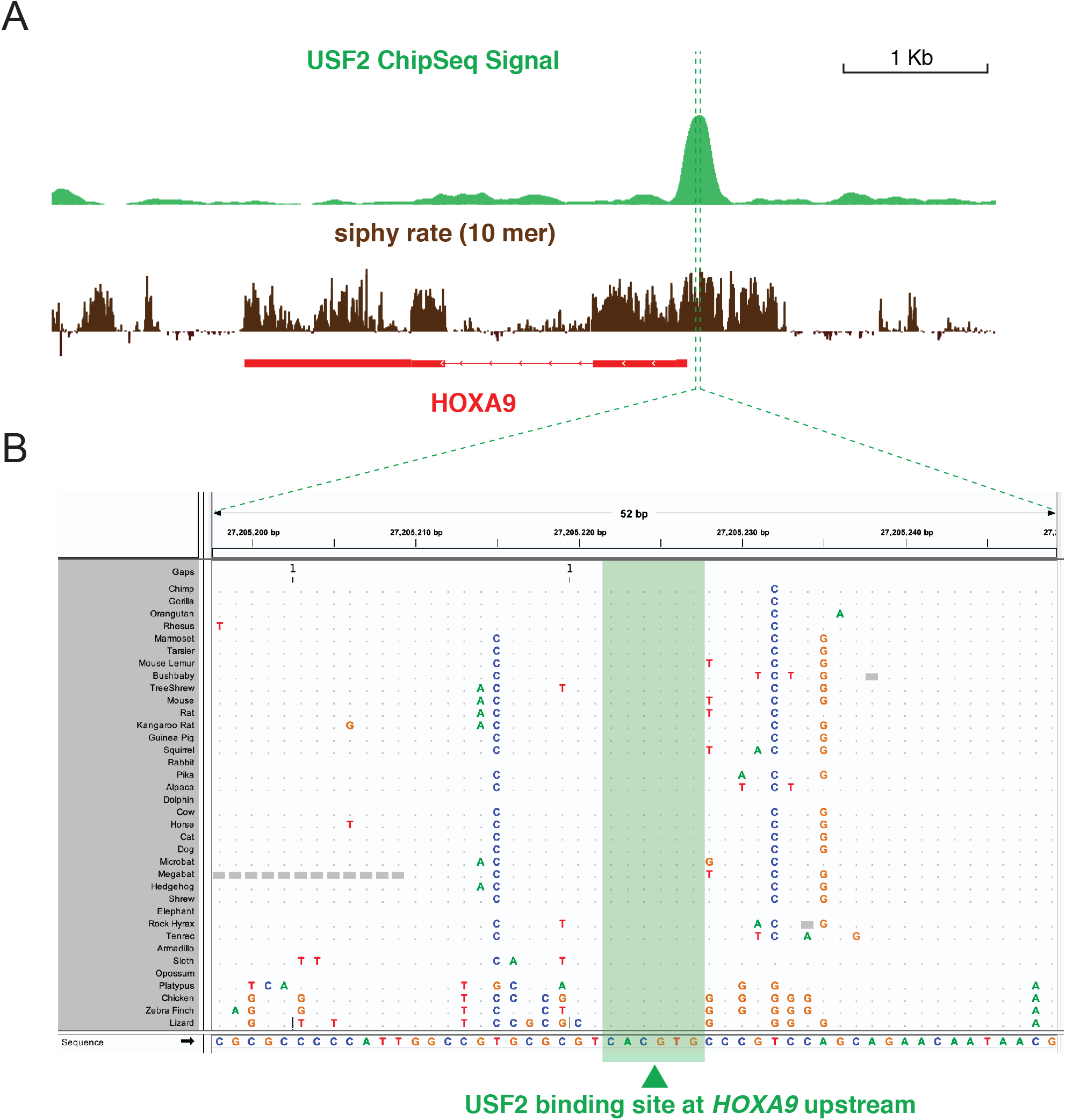
USF2 binding occupancy and transcriptional association with *HOXA9*. (A) ChIP-seq tracks from the publicly available ENCODE dataset of human ES cells identified the specific occupancy at *HOXA7* and *HOXA9* promoters. Other ChIP-seq tracks from SEM cells were used to define the open chromatin status of the locus. (B) Sequence conservation analysis of USF2 bound E-box motif (5’CACGTG3’) among different species.

**Figure S8.**
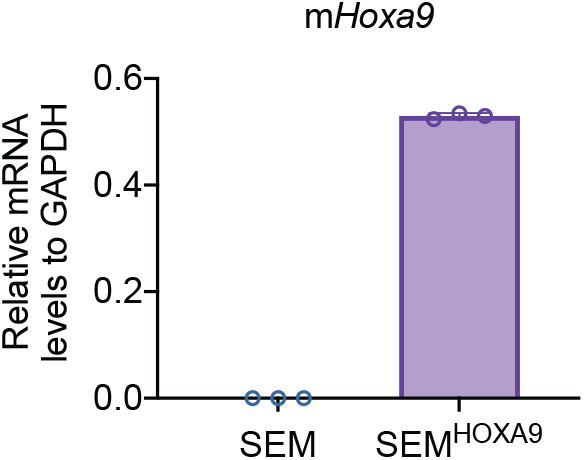
Validation of ectopic overexpression of HOXA9. A retroviral mouse *Hoxa9* expression cassette was infected into SEM cells followed by quantification of Q-PCR using specific primers against mouse *Hoxa9* coding sequence. Data are means ± SEM from two independent experiments.

**Figure S9.**
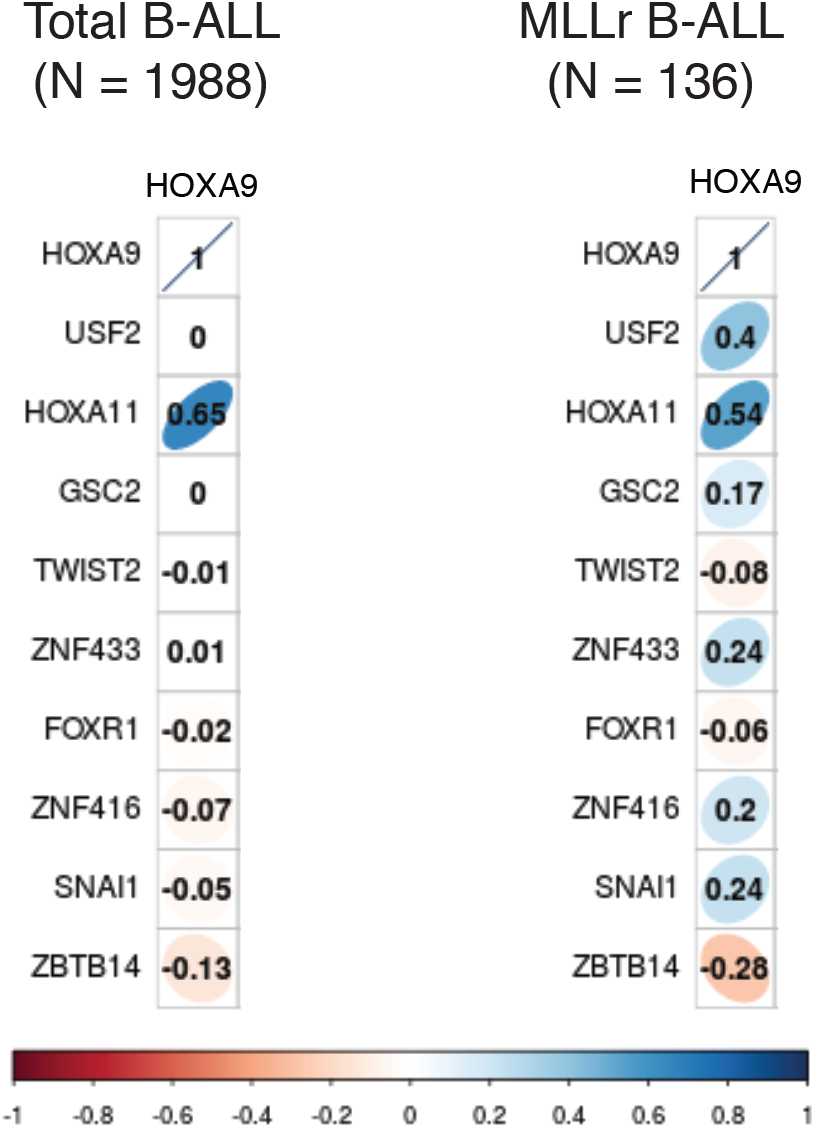
Transcriptional correlation between *USF2* and *HOXA9* in patient cohorts. Pearson’s correlation of transcriptional levels of *HOXA9* and top 10 positive regulators identified from TF screen in a cohort of 1,988 B-ALL patients (26).

**Figure S10.**
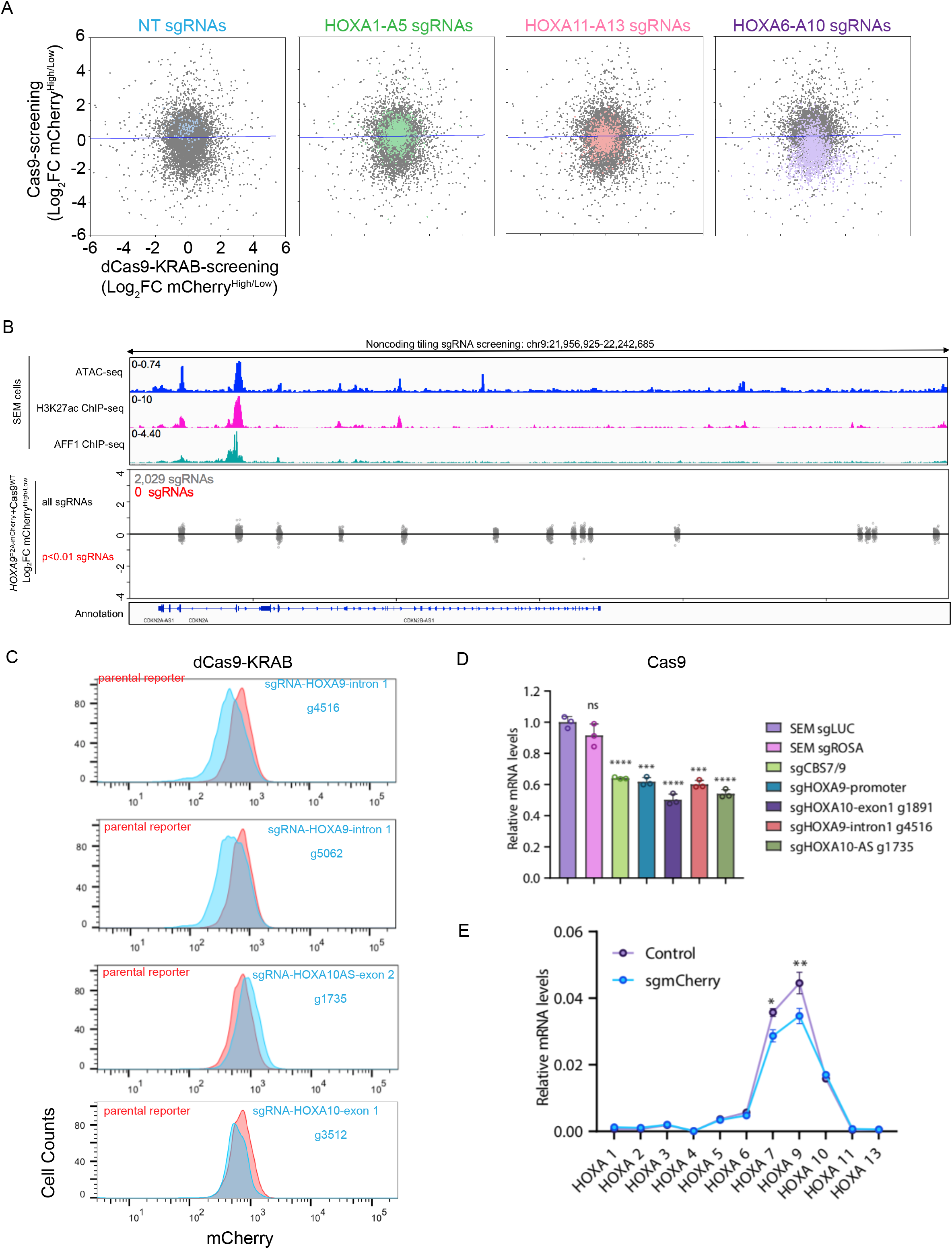
Non-coding regulation profiling of *HOXA9*. (A) The global correlation of sgRNAs enrichment in selected regions from dCas9-KRAB and Cas9 screens in the *HOXA9^P2A-Cherry^* reporter cell line. (B) The global distribution of 2,029 sgRNAs targeting the *INK4*/*ARF* locus region from the Cas9 screen in the *HOXA9^P2A-Cherry^* reporter cell line. (C) Flow cytometry analysis was conducted to validate the transcriptional regulation of *HOXA9* by screen identified *cis*-acting regulatory elements in dCas9-KRAB-expressing *HOXA9^P2A-mCherry^* reporter SEM cells infected with individual sgRNAs. (D) Q-PCR was performed to validate the transcriptional regulation of *HOXA9* by sgRNAs identified from Cas9 mediated non-coding screen. Individual sgRNAs were infected into SEM cells expressing Cas9 followed by Q-PCR assay against *HOXA9*. (E) A sgRNA against coding sequence of mCherry was infected into Cas9-expressing *HOXA9^P2A-mCherry^* reporter SEM cells followed by Q-PCR assay against *HOXA9*.

## Notes

### Competing Interest Statement

The authors have declared no competing interest.

